# Developing a Toolbox of Antibodies Validated for Array Tomography-Based Imaging of Brain Synapses

**DOI:** 10.1101/2023.06.28.546920

**Authors:** Kristina D. Micheva, Belvin Gong, Forrest Collman, Richard J. Weinberg, Stephen J. Smith, James S. Trimmer, Karl D. Murray

## Abstract

Antibody-based imaging techniques rely on reagents whose performance may be application-specific. Because commercial antibodies are validated for only a few purposes, users interested in other applications may have to perform extensive in-house antibody testing. Here we present a novel application-specific proxy screening step to efficiently identify candidate antibodies for array tomography (AT), a serial section volume microscopy technique for high-dimensional quantitative analysis of the cellular proteome. To identify antibodies suitable for AT-based analysis of synapses in mammalian brain, we introduce a heterologous cell-based assay that simulates characteristic features of AT, such as chemical fixation and resin embedding that are likely to influence antibody binding. The assay was included into an initial screening strategy to generate monoclonal antibodies that can be used for AT. This approach simplifies the screening of candidate antibodies and has high predictive value for identifying antibodies suitable for AT analyses. In addition, we have created a comprehensive database of AT-validated antibodies with a neuroscience focus and show that these antibodies have a high likelihood of success for postembedding applications in general, including immunogold electron microscopy. The generation of a large and growing toolbox of AT-compatible antibodies will further enhance the value of this imaging technique.

**Significance Statement:** Array tomography (AT) is a powerful volume microscopy technique for high-dimensional analysis of complex protein populations in cells and organelles, including synapses. AT involves the use of ultrathin serial sections embedded in resin and subjected to multiple rounds of immunofluorescence antibody (Ab) labeling and imaging. AT relies on antibody-based detection of proteins but because commercial antibodies are typically validated for other applications they often fail for AT. To identify antibodies with high probability of success in AT we developed a novel screening strategy and used this to create a comprehensive database of AT-validated antibodies for neuroscience.

## Introduction

Array tomography (AT) is a powerful technique for the analysis of large populations of synapses with deep proteomic dimensionality. AT involves preparing ultrathin serial sections from brain tissue that has been embedded in acrylic resin, and subjecting this array of sections to multiplex immunofluorescence antibody (Ab) labeling and imaging, followed by multiple rounds of iterative Ab removal, reprobing and imaging (Micheva and Smith, 2007). After many rounds of imaging, sections can be exposed to heavy metal stains, and further imaged with scanning electron microscopy. Ultimately, images are reconstructed into three-dimensional volumes of brain ultrastructure with fluorescent labeling overlays (Collman et al., 2015). This technique can simultaneously interrogate the proteomic composition of thousands of synapses with deep dimensionality (Micheva et al., 2010a; O’Rourke et al., 2012; Holderith et al., 2020). Unfortunately, many commercial Abs do not exhibit efficacy and/or specificity when applied to brain samples prepared for AT (Micheva and Smith, 2007; Micheva et al., 2010a), hindering efforts to broadly implement this powerful imaging technique.

While further refinement of tissue preparation for AT could potentially lead to improved labeling with existing antibodies, such efforts are severely limited by two considerations. First, because multiplexing is a major advantage of the method, one needs to find conditions that will be beneficial for all antibodies. Often, changing one parameter (e.g., less fixation) may improve the performance of an antibody, while decreasing the performance of other antibodies, or resulting in loss of smaller cytosolic antigens and thus hindering their detection. Second, the ability to preserve ultrastructure and use both immunofluorescence and electron microscopy is a key feature of AT. Antigenicity can be improved by resin removal (e.g. Holderith et al. 2020, Cell Rep. 32:107968), but this damages the ultrastructure making it difficult to examine the tissue under the electron microscope (Brorson, 2001). Therefore, we focused our efforts on generating and validating a set of Abs with high efficacy and specificity for brain tissue prepared using current AT protocols.

We had previously developed a reliable pipeline for generating, screening and validating monoclonal antibodies (mAbs) for neuroscience research, initially focusing on voltage-gated potassium channels (Bekele-Arcuri et al., 1996). This approach comprised analyses of numerous candidate mAbs in immunoblot and immunohistochemistry assays against mammalian brain samples (Bekele-Arcuri et al., 1996). This system reliably yielded mAbs against other ion channels (Boiko et al., 2001), synaptic scaffolds (Tiffany et al., 2000; Rasband et al., 2002), adhesion molecules (Rasband et al., 2001; Rasband and Trimmer, 2001) neurotransmitter receptors (Perez-Otano et al., 2001), and a variety of other targets. This approach was used to provide highly validated mAbs to the research community in an NIH-funded effort at the UC Davis/NIH NeuroMab Facility (Rhodes and Trimmer, 2006; Gong et al., 2016). A key aspect of Ab validation is to test them for efficacy and specificity directly in the particular application, sample type and under the exact sample preparation and labeling conditions in which they will be subsequently used (Rhodes and Trimmer, 2006; Lorincz and Nusser, 2008; Bordeaux et al., 2010; Gong et al., 2016). When a new immunolabeling technique like AT is introduced to the scientific community, it remains uncertain whether existing Ab collections will be effective and specific in the new application. Initial tests on a set of commercial Abs suggested that only a restricted subset of Abs screened on conventional assays would show efficacy and specificity for ultrathin sections embedded in plastic. Accordingly, identifying which Abs can be used on AT samples for systematic evaluation of brain synapses remains a requirement for broad and effective use of this powerful technique.

Here we describe efforts aimed at developing a reliable platform for validating Abs for AT. We present the results of using this platform in analyses of existing mAbs developed and/or validated for other purposes, and in new projects specifically aimed at developing novel mAbs for use in AT.

## Materials and Methods

### Hybridoma generation and conventional mAb screen

Mouse immunizations, splenocyte isolation, hybridoma generation and conventional screening were performed following the protocols in Bekele-Arcuri et al., 1996; and Gong et al., 2016, except that electrofusion was used to generate hybridomas. Two ELISA assays, one against purified protein immunogen, and one against transfected heterologous cells overexpressing the full-length target protein, were used in parallel as the primary screen (Gong et al., 2016). A selected set of ELISA-positive candidates were taken through subsequent screens, including immunocytochemistry on transfected cells, immunoblots on brain subcellular fractions, and immunohistochemistry on conventionally prepared brain sections (Bekele-Arcuri et al., 1996; Rhodes and Trimmer, 2008; Gong et al., 2016).

### Preparation of cell pellet arrays for AT cell-based proxy screen

The cell-based proxy screen (CBS) was developed from a previously reported protocol for preparing cultured cells for transmission electron microscopy (Schrand et al., 2010). Briefly, COS-1 cells were cultured overnight in 10 cm tissue culture plates until a confluency of ∼70% and then transfected with mammalian expression plasmids using Lipofectamine 2000 (ThermoFisher, Cat# 11668030) per manufacturer’s instructions. Cells were either co-transfected with plasmids encoding enhanced green fluorescent protein (EGFP) and the target protein of interest, or with plasmid encoding the target protein fused to a reporter tag (EGFP, FLAG). Transfected cells were incubated at 37°C/5% CO_2_ for 72 hours, then harvested in Versene with manual pipetting to release adherent cells. Cells from multiple culture plates were pooled into a single 15 mL tube and centrifuged at 1,000 x g for 5 min at room temperature (RT). The subsequent pellet was transferred to a glass vial and fixed for 2 h at RT in AT fixative (4% formaldehyde (FA) in 10 mM phosphate buffered saline (PBS, 138 mM NaCl, 2.7 mM KCl) with 2.5% sucrose, made fresh from 8% aqueous FA (Electron Microscopy Sciences (EMS), Cat# 157-8)). The pellet was rinsed three times for 10 min each in PBS containing 50 mM glycine, followed by dehydration using 5 min incubations in solutions of 50% ethanol (1X) and 70% ethanol (3X). The pellet was then washed twice for 5 min each in a solution of 3 parts LR White acrylic resin (hard grade, SPI supplies Cat# 2645) and 1 part 70% ethanol, and then four times for 5 min each in 100% LR White at 4 °C. The pellet was left in LR White overnight at 4 °C, then transferred to a gelatin capsule filled with LR White resin, capped, and incubated for 24 h at 55 °C. To generate semi-thin (400 nm) sections, the plastic “bullet” containing embedded cells was manually trimmed and then sectioned on an ultramicrotome (Leica, Ultracut UCT). Sections were collected using a thin metal loop, placed in single wells of a collagen-coated glass bottom 96 well plate (Corning 4582), air dried and stored in the dark at RT until screening.

### Immunolabeling and analyses of CBS proxy assay

Semi-thin (400 nm) sections in 96 well plates were rinsed in 50 mM glycine in Tris-buffered saline (TBS, 50 mM Tris, 150 mM NaCl, pH 7.6) for 5 min at RT. Glycine was removed and sections were incubated in blocking buffer (0.05% Tween-20 (EMS, Cat# 25564) and 0.1% BSA (EMS, Cat# 25557) in TBS) for 5 min at RT and then incubated in primary Ab in blocking buffer for 2 h at RT. Following three washes in TBS for 5 min each, sections were incubated in goat anti-mouse IgG secondary Ab conjugated to Alexa Fluor-594 for 30-45 min at RT. Following secondary labeling, sections were washed in TBS for 5 min each in RT. Sections were imaged using a 40X/1.2 NA objective on a Zeiss AxioObserver Z1 microscope with an AxioCam HRm digital camera controlled with Axiovision software (Zeiss, Oberkochen, Germany). Target labeling of Ab was assessed by comparing fluorescent signal in red (Alexa Fluor-594) and green (EGFP) channels for degree of colocalization (specificity of target label) and for labeling intensity. Labeling was rated on a scale of 0 (no label) to 4 (intense and complete colocalization).

### Preparation of arrays for AT from mouse neocortex

Arrays were prepared following the protocol in Micheva et al., 2010b. All animal procedures were performed in accordance with the Administrative Panel on Laboratory Animal Care at Stanford University. Briefly, pentobarbital anesthetized mice were subjected to intracardial perfusion with 4% FA in PB (0.1M phosphate buffer, pH 7.4) made fresh from powdered paraformaldehyde. Following removal of the perfusion-fixed brain, small (1 mm^3^) blocks of cerebral cortex were dissected and immediately transferred to AT fixative for 1 h at RT followed by overnight incubation at 4°C. Tissue blocks were then washed, dehydrated, and embedded in LR White resin according to steps described above for CBS pellets. After embedding, blocks were manually trimmed, and serial sections (70 nm) were cut with an ultramicrotome (Leica, Ultracut UCT) and collected onto gelatin-coated glass coverslips. Sections were air dried and stored in the dark at RT until ready to be labelled.

### Human neocortical tissue preparation for AT

Fresh human tissue and autopsy tissue prepared for AT from previous studies was used (Micheva et al., 2018). Briefly, human cortical samples were rinsed in saline and placed in RT fixative (4% FA in PB) for 1 h. The tissue was further fixed for 23 h at 4°C, for a total time of 24 h in fixative. The tissue was then transferred to phosphate buffered saline (PBS) with 0.01% sodium azide and stored at 4°C before further processing. The tissue was dehydrated and embedded following the same protocol as for the CBS pellet, except that incubation times in the different ethanol solutions and in resin were increased to 10 min.

### Lowicryl HM20 embedding for conjugate immunofluorescence-SEM AT applications

All animal procedures were performed in accordance with the University of North Carolina animal care committee’s regulations. After deep anesthesia with pentobarbital, adult mice (3 to 4 months old) were perfusion-fixed with a mixture of 2% glutaraldehyde/2% FA, dissolved in 0.1 M phosphate buffer (pH 6.8). Brains were removed and postfixed overnight at 4°C in the same fixative. Following extensive washes in buffer, 200 μm-thick Vibratome sections were collected, incubated on ice on a shaker with 0.1% CaCl_2_ in 0.1 M sodium acetate for 1 h, then cryoprotected through 10% and 20% glycerol, and overnight in 30% glycerol in sodium acetate solution. The next day, small tissue chunks from neocortex were dissected out and quick-frozen in a dry ice/ethanol bath. Freeze-substitution was performed using a Leica AFS instrument with several rinses in cold methanol followed by substitution in a 2–4% solution of uranyl acetate in methanol, all at -90°C. After 30 h incubation, the solution was slowly warmed to -45°C and infiltrated with Lowicryl HM-20 over 2 days. Capsules containing tissue chunks were then exposed to UV and gradually warmed to 0°C. Polymerized capsules were removed from the AFS apparatus and further exposed to UV at RT for an additional day, to complete curing of the plastic.

### Immunofluorescent labeling and analysis of brain AT sections

Ultrathin (70 nm) sections on coverslips were incubated in 50 mM glycine in TBS for 5 min at RT, followed by blocking solution (0.05% Tween 20 and 0.1% BSA in TBS) for 5 min at RT, and then incubated in primary Abs diluted in blocking solution overnight at 4°C. The reference antibodies are listed in Table 1. Following three washes in TBS for 5 min each, sections were incubated in cross-adsorbed Alexa Fluor dye-conjugated goat secondary Abs (ThermoFisher Scientific), diluted 1:150 in blocking solution for 30 min at RT. The mAbs were detected using Alexa Fluor-594 conjugated goat anti-mouse IgG (H+L) (Invitrogen Cat# A-11032) and the reference Ab with an Alexa Fluor-488 conjugated goat Ab against the appropriate host (Invitrogen, Cat# A-11034 anti-rabbit, Cat# A-11073 anti-guinea pig or Cat# A-11039 anti-chicken). Subsequently, labelled sections were washed 3 times in TBS for 5 min each, followed by three rinses in water for 30 seconds each. Coverslips with sections were mounted onto glass slides using SlowFade Gold Antifade mountant with DAPI (Invitrogen #S36964) and imaged the same day using a 63X/1.4 Plan-Apochromat 1.4 NA oil objective on a Zeiss AxioImager Z1 microscope with an AxioCam HR digital camera controlled with Axiovision software (Zeiss, Oberkochen, Germany). Image ZVI files were converted to TIFF and uploaded into Fiji Imaging software for analysis. Images of multiplex labeling from at least three serial sections were aligned using the DAPI signal with the MultiStackReg plugin in FIJI (Thevenaz et al., 1998; Schindelin et al., 2012) and immunolabeling was assessed for proper localization against a reference marker. Quality of labeling was assessed by experienced observers and rated on a scale of 0 (no label or off target only) to 4 (target only label).

**Table 1.**
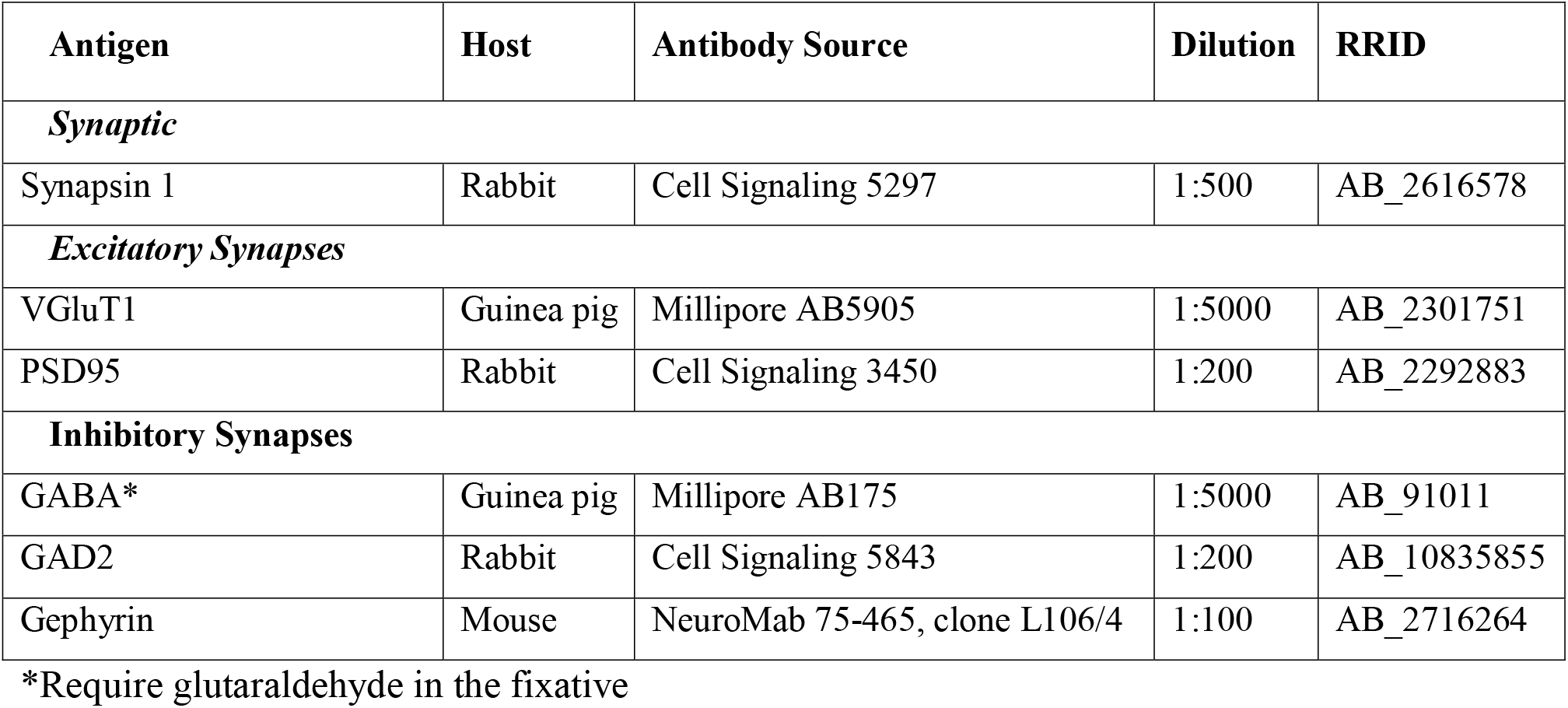
List of reference antibodies used for antibody screening. RRID (Research Resource Identifier). *Requires glutaraldehyde in the fixative

### Synaptic Antibody Characterization Tool (SACT) Analysis

To identify top Ab candidates for synaptic target localization we used the SACT program (Simhal et al., 2018), which applies an unsupervised probabilistic detection algorithm (Simhal et al., 2017) to identify fluorescent puncta and determine whether they are located at synapses. For each candidate Ab, the size, volume and density of immunolabeled puncta was measured and compared with similar measures made using an AT synaptic marker reference Ab in the same sections (synapsin 1 or PSD-95). To evaluate the sensitivity and specificity of a candidate Ab we plotted the target synaptic density of each candidate (defined as the number of synapses detected with the candidate Ab per unit volume) and the Target Specificity Ratio (TSR), defined as the number of synapses detected by the candidate Ab relative to the total number of Ab puncta (Simhal et al., 2018).

### Immunogold labeling of osmium-treated samples

For immunogold EM of osmium-treated tissue embedded in LR White, the samples were prepared similarly to Immunofluorescence AT, except that the fixative contained 0.1% glutaraldehyde in addition to the 4% FA, and a postfixation step was added with osmium tetroxide (0.1%) and potassium ferricyanide (1.5%) with rapid microwave irradiation (PELCO 3451 laboratory microwave system with ColdSpot; Ted Pella, Redding CA), 3 cycles of 1 min on-1 min off-1 min on at 100 W, followed by 30 min at RT. The immunolabeling protocol was similar to the immunofluorescence labeling, with two additional steps in the beginning: treatment for 1 min with saturated sodium metaperiodate solution in dH_2_O (to remove osmium) and 5 min with 1% sodium borohydride in Tris buffer to reduce free aldehydes resulting from the presence of glutaraldehyde in the fixative. A 10 nm gold-labeled goat anti-mouse IgG secondary Ab (SPI Supplies, West Chester, PA) was used at 1:25 for 1 hr. After washing off the secondary Ab, the sections were treated with 1% glutaraldehyde for 1 min to fix the Abs in place. The sections were post-stained with 5% uranyl acetate for 30 min and lead citrate for 1 min.

### Immunogold labeling of Lowicryl HM20-embedded tissue

Thin sections (∼80 nm) of adult mouse cortex embedded in Lowicryl HM20 were cut and collected on nickel mesh grids. Grids were blocked in 1% bovine serum albumin in Tris-buffered saline pH 7.6 with 0.005% Tergitol NP-10, and incubated overnight at 21–24°C with the primary Ab. Grids were then rinsed, blocked in 1% normal goat serum in Tris-buffered saline pH 8.2, and incubated in goat anti-mouse secondary Abs conjugated to 10 nm or 20 nm-diameter gold particles (Ted Pella, Redding CA). Grids were counterstained with 1% uranyl acetate, followed by Sato’s lead, and examined in a Philips Tecnai transmission electron microscope at 80 KV, and images collected with a 1024 × 1024 cooled CCD (Gatan).

## Results

### Efficacy and specificity of commercial Abs against synaptic proteins for array tomography

Initial efforts to identify AT-appropriate Abs that successfully labelled target proteins in sections from FA-fixed and LR White embedded mouse neocortex relied on ad hoc sampling of the vast array of pre-existing Abs from commercial sources. Over 300 commercial Abs, selected based on available literature and personal communication, were evaluated for efficacy and specificity for AT (**Supplemental Table 1**). Criteria for success included immunolabeling that matched known cellular expression and subcellular localization of the target protein. When synaptic proteins were targeted, the subcellular distribution of the immunofluorescence of the tested Ab was evaluated by assessing colocalization with well-known reference synaptic Abs (**Figure 1**, **Table 1**) and other AT validated antibodies (Supplemental Table 2). All of the reference antibodies were characterized extensively and used in several previously-published studies (e.g. Micheva et al. 2010a, Collman et al. 2015; Simhal et al. 2018). Potential background or nonspecific labeling was evaluated using “exclusion” markers defined for each target protein; for example, inhibitory synapse markers when testing Abs against proteins thought to be restricted to excitatory synapses. The labeling pattern of each Ab was compared with that for the nuclear marker DAPI to control for background nuclear immunolabeling (**Figure 1**). Scoring was performed using visual inspection of images by a trained observer. We found that even with widely-used commercial antibodies generally considered to yield optimal results, up to 50% fail completely (139/306 tested, Supplemental Table 1). Even more alarming was the observation that for 32% of the targets (63 out of 196) we failed to identify an AT-suitable commercial antibody. Therefore, we set out to design a more focused and application-specific screening process.

**Figure 1.**
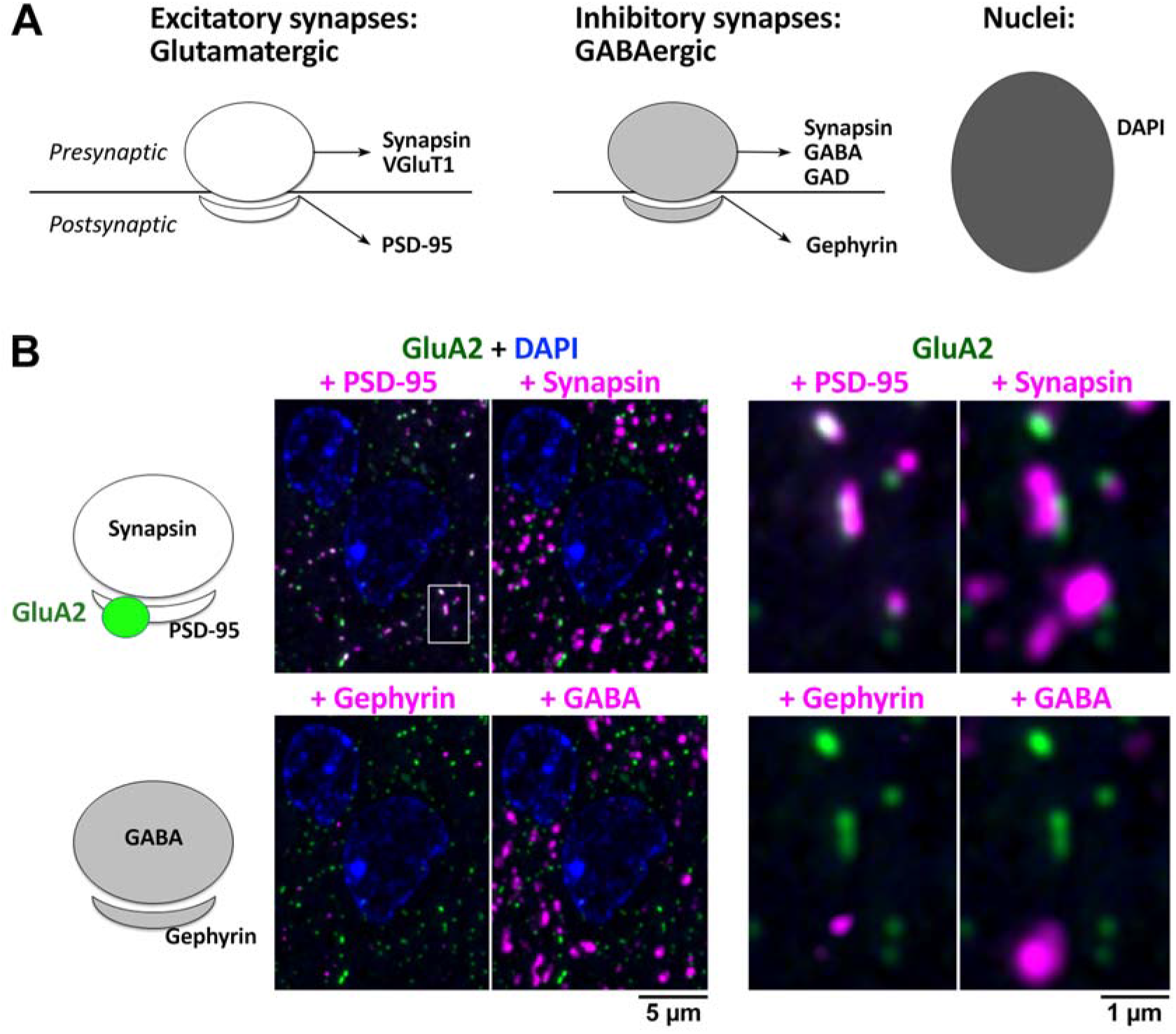
Initial AT evaluation strategy for identifying synaptic Abs validated in other applications. **A**. Common reference markers for pre-and postsynaptic locations and nuclei. **B**. Example evaluation of an Ab against GluA2 (Abcam ab206293), a glutamatergic receptor with a known postsynaptic localization at excitatory synapses. A single 70 nm section from adult mouse cortex, labeled with the GluA2 Ab (green) and synaptic markers PSD-95 (Cell Signaling 3450), synapsin (Synaptic Systems 106006), gephyrin (NeuroMab L106/93), GABA (Millipore AB175), and the nuclear label DAPI. The panel to the right is an enlarged view of the boxed area in the left panel. The GluA2 Ab was scored as excellent, based on its colocalization with PSD- 95, adjacency to synapsin and minimal background label. See Supplementary Tables 1 and 2 for more details.

### Finding application-specific anti-PSD-95 mAbs via retrospective screen of a prior monoclonal project

We previously performed a project to develop mAbs recognizing the mammalian synaptic marker PSD-95 employing a region of human PSD-95 (amino acids 77-299 of Uniprot accession number P78352-2) as the immunogen. This resulted in a set of 96 independent samples that displayed immunoreactivity against a recombinant PSD-95 protein fragment by ELISA, and to varying degrees on immunoblots and by immunohistochemistry against brain samples (Tiffany et al., 2000; Rasband et al., 2002). From these 96 samples we selected one mAb, K28/43, that exhibited efficacy and specificity for reliable labeling of mammalian PSD-95 in brain tissue sections and cultured neurons. However, all 96 ELISA-positive samples had been archived as frozen hybridomas for potential future use.

In a subsequent analysis of mAbs from this project aimed at identifying mAbs recognizing Zebrafish PSD-95, we observed that binding to mammalian PSD-95 was not predictive of labeling the Zebrafish ortholog (Meyer et al., 2005). Whereas clone K28/43 robustly recognized mammalian PSD-95 (Rasband et al., 2002), it did not recognize Zebrafish PSD-95, although other mAbs from this same project did (Meyer et al., 2005). Human and Zebrafish PSD-95 (Uniprot accession number A0A8M3ASX4) share 89% amino acid identity within the region used as the immunogen, with distinct regions of high and low sequence identity. This provides a likely basis for mAbs with distinct epitopes within the collection originally selected for their binding to human PSD-95 displaying differences in binding to Zebrafish PSD-95. That we were successful in rescreening existing mAbs within this collection for a new purpose suggested that this would be a viable approach to identify mAbs for not only new targets, but also for new applications, without the need to launch new mAb projects from scratch.

Accordingly, we evaluated mAbs for labeling of PSD-95 in samples processed for AT (**Figure 2A**). Following the strategy outlined in **Figure 1**, controls to assess specificity included co-labeling with different reference Abs against the same target (a rabbit monoclonal anti-PSD- 95 Ab, Cell Signaling #3450) and Abs against the adjacent presynaptic compartment of excitatory synapses (a rabbit monoclonal anti-synapsin Ab, Cell Signaling #5297). Because postsynaptic densities of synapses usually span at least two adjacent ultrathin sections (70 nm thickness each), the consistency of labeling was further assessed by examining serial sections for the presence of immunolabel. Using these criteria, we observed robust and specific labeling with K28/43 of excitatory synapses in AT sections prepared from mouse cortex, and from freshly obtained resected human neocortex (**Figure 2B**). However, no specific labeling with K28/43 could be detected in neocortical samples from human autopsy brain, even though the tissue was fixed and embedded using the same protocol. Therefore, we expanded our search to include other mAbs from the K28 anti-PSD-95 project that like K28/43 had been identified as labeling brain tissue prepared using conventional immunohistochemistry protocols. We found that unlike K28/43, mAbs K28/37, K28/74 and K28/77 labeled synapses in both fresh and autopsy human brain samples, as well as in mouse neocortex samples.

**Figure 2.**
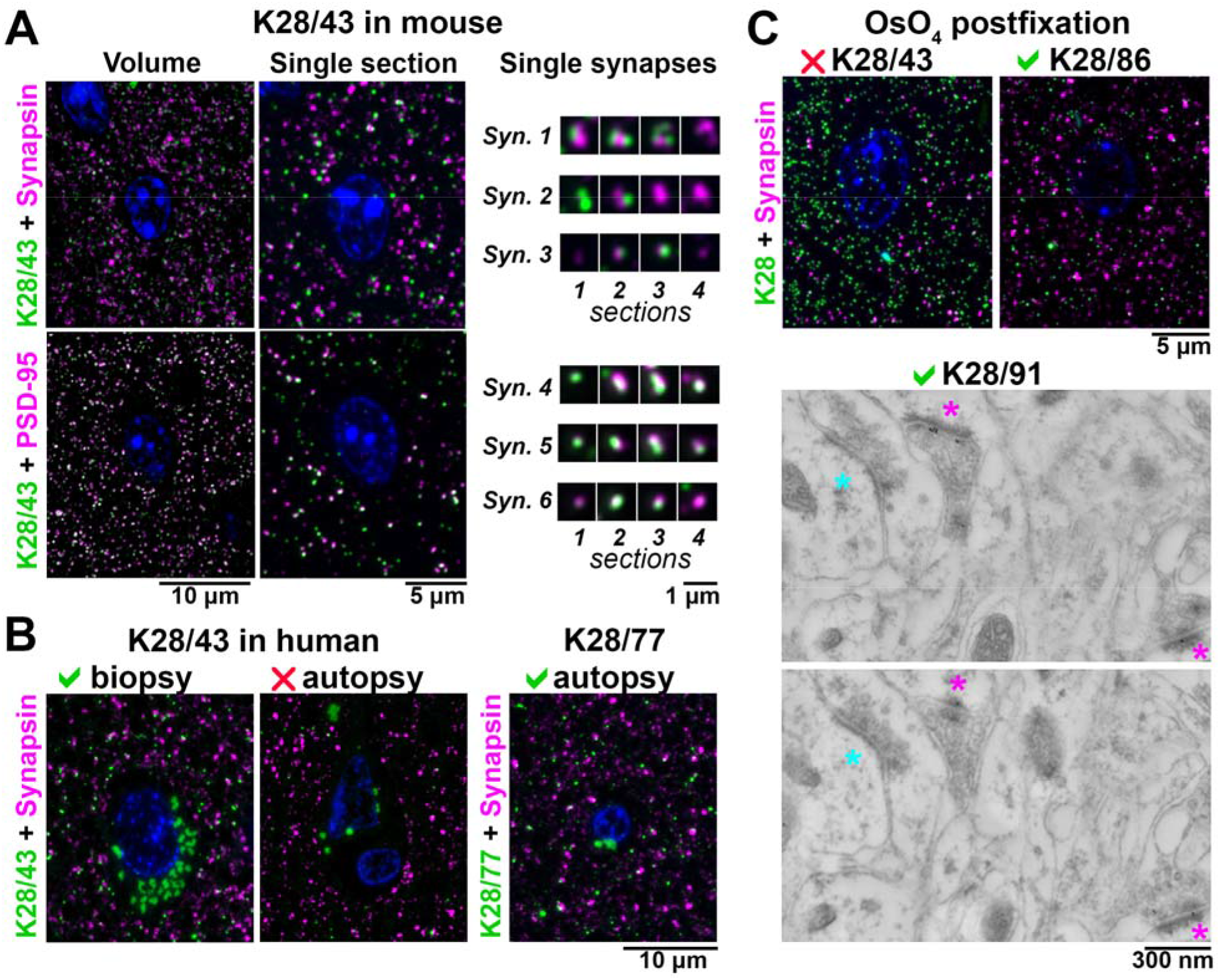
Application-specific performance of mAbs. **A**. Ultrathin sections from LR White-embedded mouse neocortex immunolabeled with anti-PSD-95 mAb K28/43 (green) and a reference anti-synapsin Ab (Cell Signaling #5297, magenta, top), or a reference anti-PSD-95 mAb (Cell Signaling ##3450, magenta, bottom). Nuclei are labeled with DAPI (blue). To the right, examples of individual synapses are shown, with four serial sections through each. Synapses 1 – 3 are immunolabeled with K28/43 (green) and anti-synapsin Ab (magenta), and synapses 4 – 6 with K28/43 (green) and the reference anti-PSD-95 mAb (magenta). **B**. Immunolabeling of human neocortical samples from biopsy or autopsy with the same K28/43 mAb. While K28/43 performs well on human biopsy tissue (left), it shows very sparse labelling on autopsy tissue (middle). However, a different mAb from the same project, K28/77, gives a specific and robust signal on human autopsy tissue (right). Autofluorescent lipofuscin granules, which are much more abundant in the human tissue are seen in the green channel, within the neuronal cytoplasm surrounding the nuclei. **C top.** Mouse neocortex postfixed with osmium tetroxide and immunolabeled with an anti-PSD-95 mAb (green) and a reference anti-synapsin Ab (Cell Signaling #5297, magenta). K28/43 gives dense non-specific label, but mAb K28/86 from the same project performs well in this preparation. **C bottom.** Immunogold electron microscopy of mouse neocortex with K28/91, two serial sections are shown. Excitatory synapses, recognized by their asymmetric synaptic junction (magenta asterisk), have associated immunogold particles, whereas inhibitory synapses (cyan asterisk, symmetric synaptic junction) do not.

Variations in the preparation of the AT samples also affected the performance of the mAbs. Thus, while K28/43 labeled conventional mouse neocortex AT sections, it did not label AT sections after tissue treatment with osmium (**Figure 2C**), a preparation condition commonly used to preserve ultrastructure and provide contrast for EM. However, mAbs K28/38, K28/74, K28/86 and K28/91 labeled osmium-treated tissue in AT sections, and also subsequently yielded specific immunogold labeling (**Figure 2C**). Overall, immunogold labeling on osmium-treated tissue was not very efficient and usually resulted in low labeling density, prompting us to use a different method for tissue preparation for EM purposes, as detailed below. Results from these *post facto* analyses of an existing collection of PSD-95 mAbs illustrated that application-specific re-evaluation of mAbs can identify those with strong and specific labeling that may not be identified in other assays. In addition, they highlighted the potential for retrospective analyses of other archived mAb projects to identify mAbs with characteristics suitable for use in AT.

### Application-specific generation and validation of mAbs for AT

Our generation of mAbs for neuroscience employs a stepwise screening workflow that incorporates the tissue culture aspects of classical hybridoma generation, expansion and archiving (immunization, hybridoma fusion, cell culture, cryopreservation) and parallel screening (ELISA, immunocytochemistry on transfected heterologous cells), while also including assays (immunoblots and immunohistochemistry) performed on mammalian brain samples (Trimmer et al., 1985; Bekele-Arcuri et al., 1996; Rhodes and Trimmer, 2006; Gong et al., 2016). The above experience in rescreening the PSD-95 mAb clones suggested that including samples prepared for AT would help identify mAbs useful for that application. To define AT-compatible mAbs, we first added an additional screen comprising immunolabeling and analysis of AT brain sections into our mAb pipeline (**Figure 3**). However, we found that screening with AT on brain sections was too slow and labor-intensive, given the large number of samples that needed to be prepared, immunolabeled and evaluated. Moreover, developing an alternative cell-based proxy AT assay represented an opportunity to reduce the need for animal tissues. We therefore developed a rapid and straightforward cell-based proxy assay for mAbs able to recognize their target in samples prepared as for AT, employing transiently transfected cells as used in the immunocytochemistry screening step (**Figure 3**, bottom).

**Figure 3.**
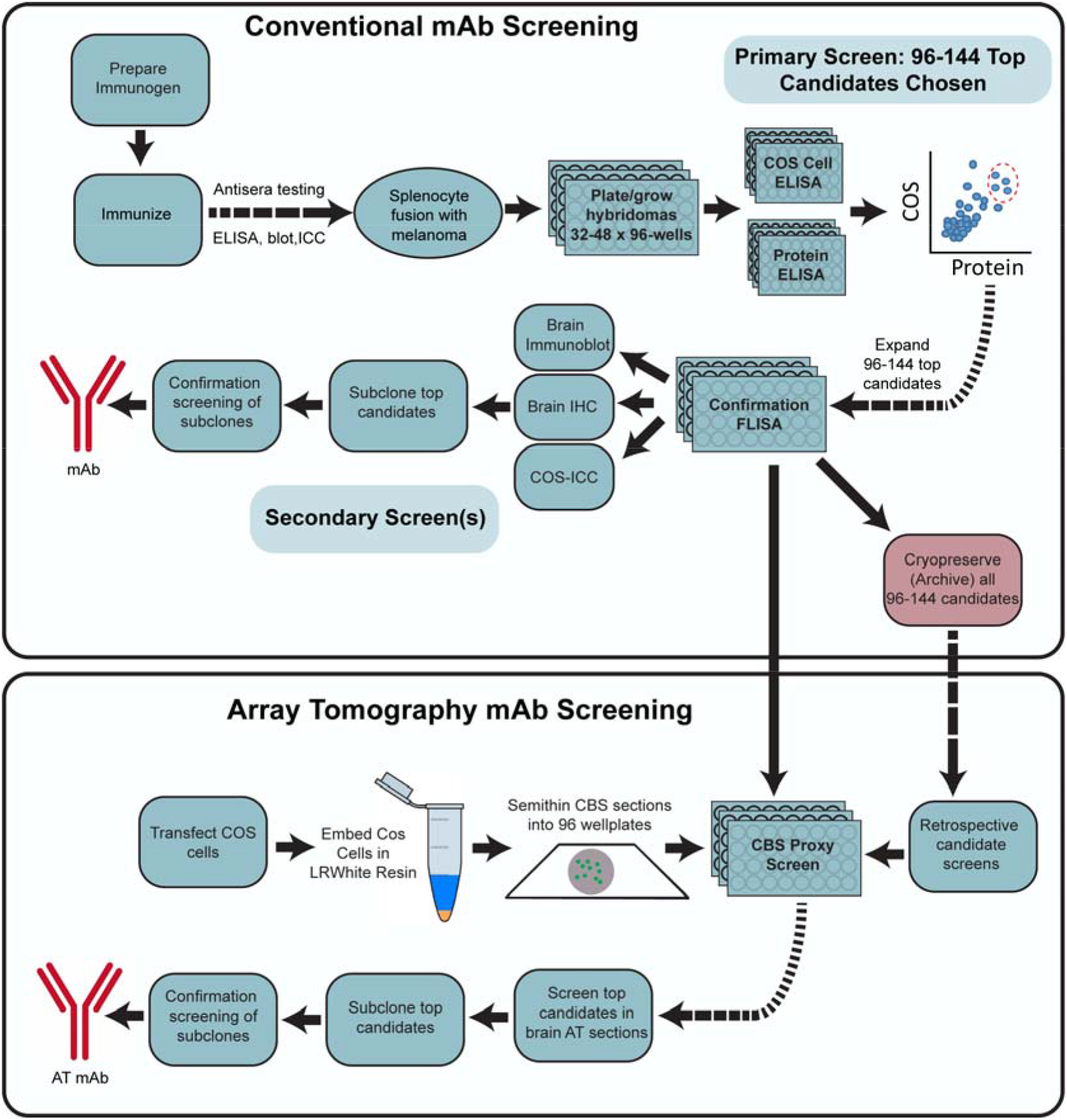
Flow diagram of the mAb screening workflow. Flow charts illustrating steps for conventional (Top) and AT-inclusive (Bottom) mAb screens.

### Homer 1L mAb generation as an exemplar mAb project

An exemplar mAb project (the L113 project) targeted Homer1L, an important component of the postsynaptic density of excitatory synapses (Xiao et al., 2000; Fagni et al., 2002; Brandstatter et al., 2004). We immunized a set of mice with a recombinant protein comprising the C-terminal two-thirds of the mouse Homer1L protein (a.a. 121-363 of accession number Q9Z2Y3-1), a primary sequence that is 97.8% identical to human Homer1L and 97.1% identical to rat Homer1L. This fragment contains an N-terminal region present in all Homer1 splice variants, and a C-terminal region unique to the longest splice variant Homer1L (Shiraishi-Yamaguchi and Furuichi, 2007). Next, we performed two sets of ELISAs on hybridoma conditioned culture medium (tissue culture supernatants; “TC supes”) harvested from individual wells of 32 x 96 well hybridoma culture plates. One set of ELISAs was against the purified fragment of Homer1L that was used to immunize the mice, and the other was against heterologous cells that had been transiently transfected to express the full-length mouse Homer1L protein and then fixed with 4% FA and permeabilized with 0.1% TX-100 (standard conditions for immunocytochemistry). We used the combined results from these two ELISAs to inform the selection of 144 hybridoma cultures for further screening, from the 2,944 samples evaluated. A scatter plot comparing results from these two distinct Homer1L ELISAs is shown in **Figure 4A**; data points for the 144 candidates selected for expansion in tissue culture and further analysis in the screening workflow are shown in light purple. Samples of TC supes harvested from the expanded cultures of these 144 selected hybridoma samples were assayed in parallel for efficacy and specificity using fluorescent immunocytochemistry against transiently transfected heterologous cells expressing full-length Homer1L (**Figure 4B**); immunoblots on brain samples (**Figure 5A**); and DAB-HRP immunohistochemistry on fixed, free floating brain sections (**Figure 5B**). The positive candidates from these assays represent distinct, partially overlapping subsets of the original 144 ELISA positive clones (**Figure 4**), with different subsets of mAbs exhibiting efficacy in each assay. In parallel, we also subjected these 144 TC supes to a novel AT-specific cell-based proxy assay as described in the following section.

**Figure 4.**
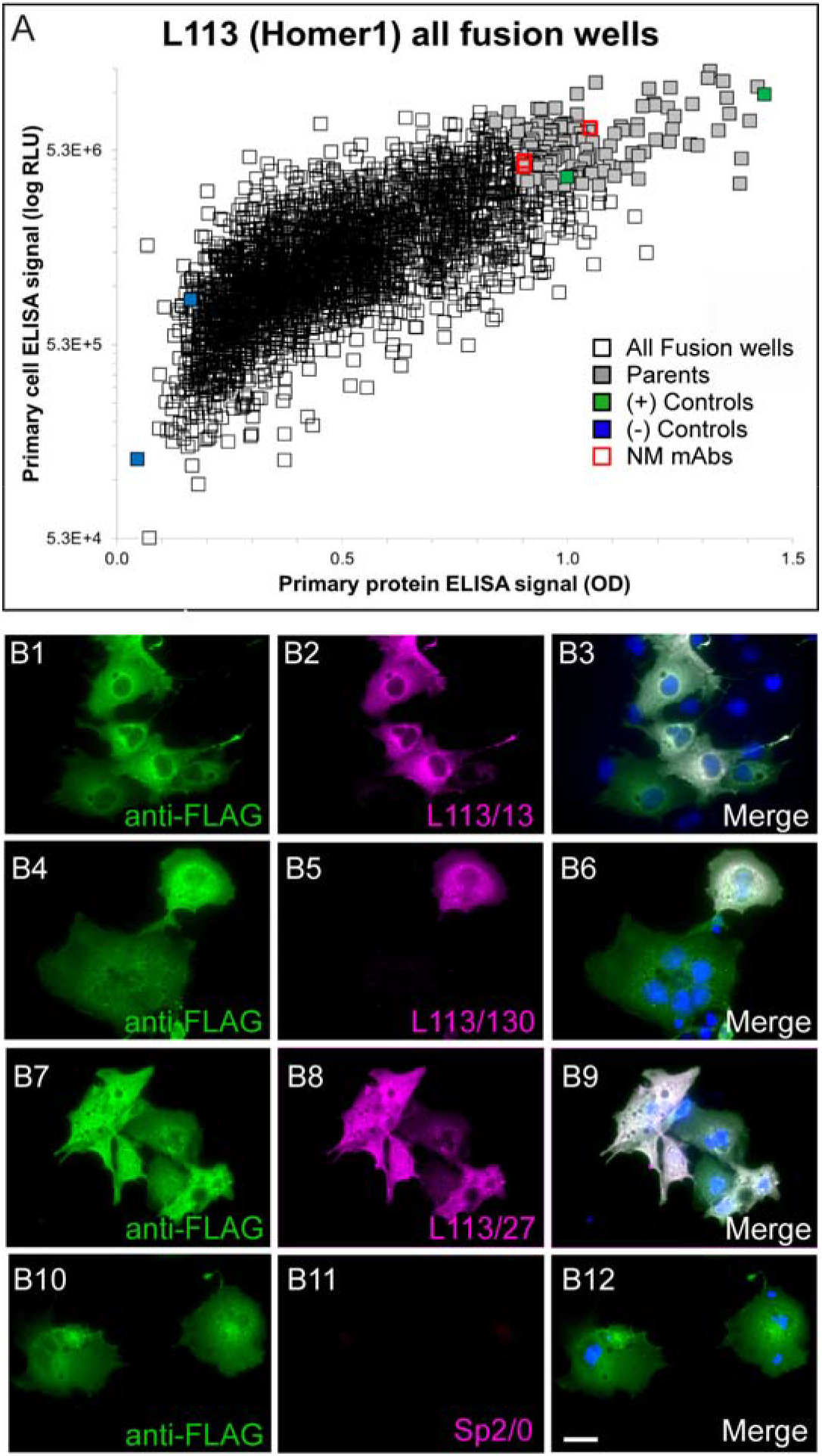
Conventional ELISA and COS-IF screening results for the L113 project targeting Homer1L. **A.** ELISA primary screen data for the project, with protein and cell ELISA data plotted on the *x*- and *y*-axes, respectively. 2,944 hybridoma samples were screened by two ELISA assays, which also included positive (green) and negative (blue) control wells. The 144 candidates selected for further screening are in grey, and the red squares denote the wells with candidates (L113/13, L133/27, L113/130) that were ultimately selected as NeuroMab mAbs. **B**. Exemplar results of the secondary COS IF screen for the L113 project. Photomicrographs show fluorescent immunolabeling of COS-1 cells transiently transfected with Flag-tagged mouse Homer1L mammalian expression construct using a rabbit anti-Flag pAb (green)(Sigma, Cat# F7425), candidate mouse <u>mAbs</u> (magenta) and Hoechst nuclear stain (blue). The first three rows show images from three positive candidates (L113/13, L113/27, L113/130 and) that were eventually selected as NeuroMabs, and the fourth row shows the negative control (Sp2/0 myeloma cell medium). Scale bar = 5 µm.

**Figure 5.**
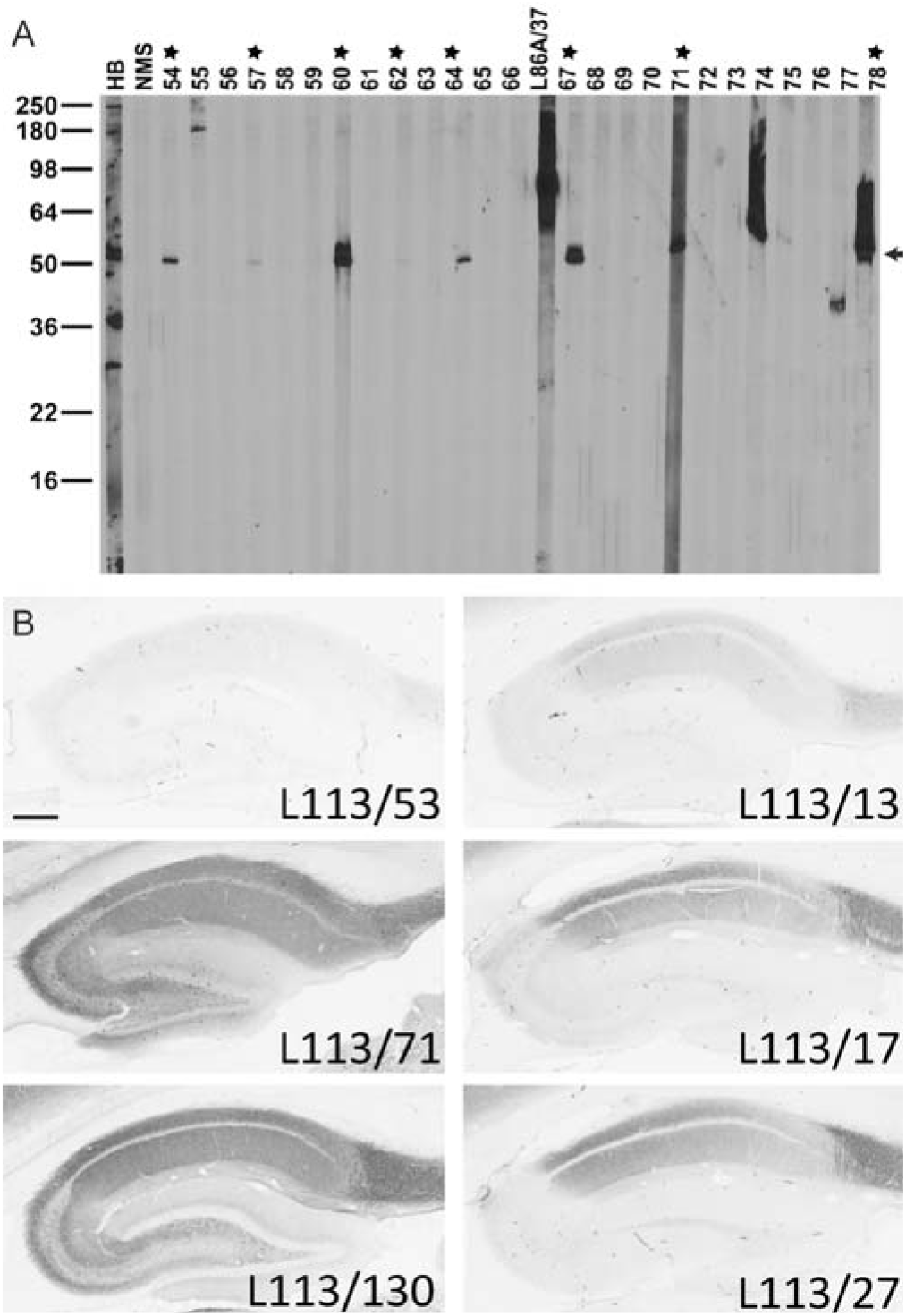
Conventional immunoblot and IHC screening results for Homer1. **A.** Representative immunoblot strips from the L113 screen. Values on left show the mobility of molecular weight standards in kDa. Each lane represents a replicate strip containing a crude rat brain membrane fraction probed with a different candidate or control Ab. Other lanes include antiserum from one of the immunized mice (HB), a mouse serum negative control, and a positive control NeuroMab mAb against a different target (L86A/37, AMIGO-1). Strips for candidates L113/54 to L113/78 are shown. Stars = positive candidates on strip blot. Arrow = expected electrophoretic mobility of Homer1L. **B.** Representative images from the L113 IHC screen. Photomicrographs show DAB/NAS immunolabeling of sagittal rat brain sections. Results from 6 candidate mAbs highlight a range of results from negative (L113/53) to partial (L113/17, L113/27, L113/13) to full (L113/71, L113/130) labeling, with the expected cellular and subcellular labeling pattern based on in situ hybridization and immunohistochemistry evidence gleaned from the literature and from publicly accessible in situ hybridization databases. Scale bar = 1 mm.

### Generating and validating mAbs for array tomography: AT-focused screening of an anti-Homer1L mAb project

Our standard mAb generation pipeline did not contain a single assay that reliably predicted which candidate mAbs would successfully label AT-prepared brain tissue (see above).

Moreover, screening all ELISA-positive clones (typically 96 or 144) on conventional AT sections of plastic-embedded brain samples, which included manual scoring by trained observers, was excessively slow and labor-intensive. This was especially problematic because we strive to identify the best candidates for subcloning of the hybridoma cell cultures to monoclonality prior to their cryopreservation, which needs to occur within one week after the initial ELISA screen (Gong et al., 2016). Therefore, we developed a novel cell-based proxy screen with a high predictive value for mAbs that would ultimately prove to be effective on labeling brain tissue in plastic-embedded AT sections. We hypothesized that the major factor distinguishing antigenicity in AT brain sections from conventional IHC is the process (dehydration, resin infiltration, heat curing) involved in the embedding of AT samples in array plastic. Our standard mAb screening workflow employs immunofluorescence labeling of the target protein expressed in heterologous cells as an important screen (**Figures 3 and 4**) (Bekele-Arcuri et al., 1996; Gong et al., 2016). By employing transiently transfected cells, the samples assayed are a mosaic population of cells with high levels of target protein expression adjacent to non-expressing cells. Since the identity of the transfected cell subpopulation is apparent by the use of an independent marker, it is easy to determine which candidates selectively label target-expressing cells. We predicted that plastic embedding of transiently transfected heterologous cells expressing target protein would provide a similarly quick, inexpensive and effective screen for candidate TC supes that exhibit target protein labeling under AT conditions.

For this cell-based screening assay, we transiently co-transfected heterologous COS-1 cells such that ≈50% of the cells were transfected to express both the target protein and an independent transfection “marker” to monitor transfection efficiency and to identify the transfected cells. Transfection markers were either encoded on separate plasmids (e.g., EGFP) cotransfected with target expression plasmids, or were encoded as tags fused to the target protein (e.g., an epitope tag or a fusion protein). For the CBS AT assay, transiently transfected cells were harvested three days post-transfection, pelleted by centrifugation, fixed in suspension, re-pelleted and the fixed cell pellet embedded in LR White plastic. A portion of the same transfection cocktail was used to transfect cover slips containing cultured COS-1 cells that were subjected to conventional immunocytochemistry, to verify successful coexpression of both the marker and the target protein. Embedding of the cell pellet in AT plastic was performed using the same protocol as used for embedding brain tissue in plastic for AT (4% FA fixation, dehydration in an ethanol series, embedding in LR White resin and curing at 55°C for 24 h). Semithin (400 nm) sections that contained a mosaic of cells overexpressing the target protein and marker adjacent to cells devoid of target protein expression (**Figure 6**) were cut and deposited into the wells of a collagen-coated clear bottom 96 well plate, which was used to screen up to 94 candidate TC supes (using the remaining two wells for positive and negative controls). The nucleus of each cell was labeled with Hoechst 33258, and bound primary Ab was detected using Alexa Fluor-594 conjugated anti-mouse IgG secondary Abs. Subsequent visual analysis was based on determining the presence of Alexa Fluor-594 signal and whether it was specific to the subset of cells with marker expression in green (**Figure 6**).

**Figure 6.**
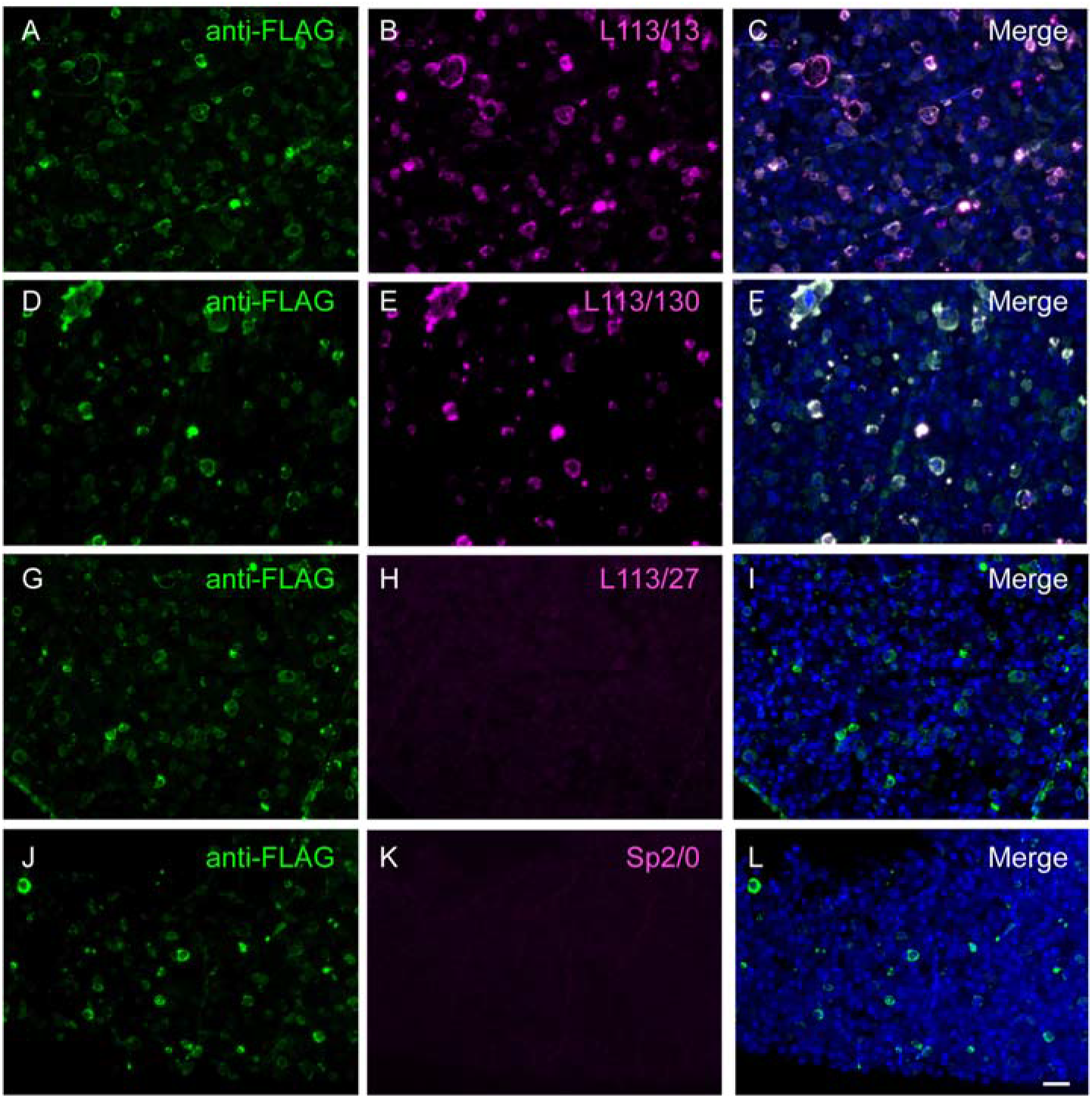
CBS assay identifies potential AT-compatible mAbs. Images of LR White embedded Homer1L-expressing transiently transfected COS-1 cells in semi-thin (400 nm) sections and labeled with candidate L113 mAbs. Only two of the three mAbs selected on the basis of their excellent performance in ELISA and conventional IHC screening were found to perform well on these AT proxy sections (L113/13, **A-C)** and L113/130 (**D-F**). mAb L113/27 does not selectively recognize the target expressing cells (**G-I**) and is similar in appearance to the negative control, conditioned medium from the Sp2/0 myeloma cell line (**J-L**). Scale bar = 50 µm.

### The AT CBS assay effectively predicts mAbs that label target proteins in AT brain sections

A proxy screen should be rapid, simple, and inexpensive, but most importantly it must be able to identify Abs that are effective when employed for their intended end use. We interrogated positive and negative samples from the AT CBS assay for labeling efficacy and specificity against brain samples in AT plastic. We found a high degree of concordance: positive samples from the AT CBS assay were much more likely to be scored as positives against AT brain sections than the population as a whole, and negatives from the CBS assays were rarely positive against AT brain sections (**Figure 7A**). For example, in the exemplar Homer1L mAb screen, of the 144 candidate mAbs screened by CBS, 60% of the CBS-positive candidates (CBS score >2) gave good brain AT labeling (brain AT score > 2.5), compared to only 4 out of 96 CBS-negative candidates (CBS score 0-1). To further assess the value of the AT CBS assay we used a previously described Synaptic Antibody Characterization Tool developed to quantitatively assess synapse Ab specificity in AT (Simhal et al., 2018). Using this tool we measured the Target Specificity Ratio (TSR) which quantifies the ratio of target Ab label (e.g., Homer1L candidate mAbs) colocalizing with a reference synaptic marker, for example Abs against synapsin 1 (Simhal et al., 2017; Simhal et al., 2018). We observed a positive correlation between TSR scores for brain AT labeling and the respective CBS scores, supporting that the CBS assay was highly predictive in identifying brain AT compatible candidate mAbs (**Figure 7B**). The target synapse density, i.e. the number of synapses detected with the antibody per unit volume reflects the sensitivity of the antibody, and was another metric used to characterize candidate antibodies. Antibodies with high brain AT score tend to have higher synapse density (Figure 7C) and higher TSR as measured with the Synaptic Antibody Characterization Tool. Together these observations illustrate that the CBS assay can identify Ab samples with a high likelihood of successful brain AT labeling.

**Figure 7.**
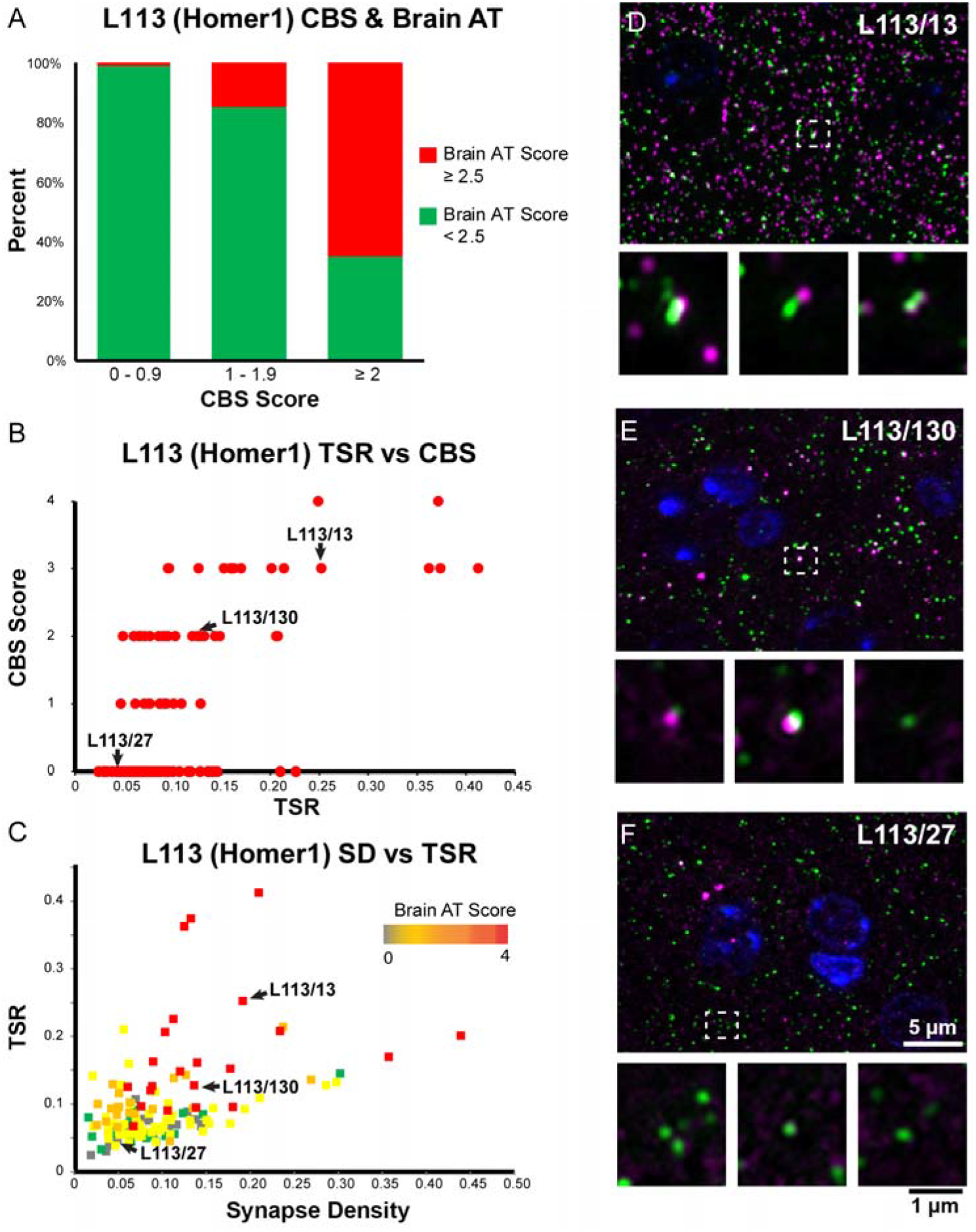
CBS positive mAbs screened on brain tissue embedded for AT. **A.** Percent of candidate mAbs with high brain AT score among mAbs with different CBS scores. Candidate mAbs with low CBS scores (0-1 and 1-2) are very unlikely to have a high brain AT score, while the majority of candidate mAbs with high CBS scores also scored high on brain AT. **B**. Correlation between TSR scores which measure Ab specificity in AT brain labeling, and CBS scores. **C**. Target synapse density which measures the Ab sensitivity in AT brain labeling plotted against the TSR scores. **D – F.** Images of ultrathin sections from LR White-embedded mouse neocortex immunolabeled with the Homer1L mAbs (magenta) L113/13 (**D**), L113/130 (**E**) (both CBS positive) and the CBS negative L113/27 (**F**), double labeled with a PSD95 Ab (green). Nuclei are labeled with DAPI (blue). The bottom of each panel includes examples of individual synapses with three serial sections through each of the AT samples. Similar to their performance in the CBS assay (Figure 5), mAbs L113/13 and L113/130 show specific labeling on AT brain sections, while L113/27 does not detect the target protein.

### Using the AT CBS assay to screen Abs for immunoelectron microscopy

During our initial screening of available polyclonal and monoclonal Abs against synaptic proteins for AT we had also observed that a high proportion of Abs effective for immunofluorescence AT on LR White-embedded sections also perform well on Lowicryl HM20-embedded sections (82%, 73 out of 89 Abs). Because Lowicryl embedding provides EM ultrastructure superior to that of LR White, we wondered whether the CBS AT assay could also screen for Abs effective for postembedding immunogold EM of brain samples in Lowicryl. We first tested whether the mAbs identified in the AT CBS screen performed on cells embedded in LR White plastic would also exhibit effective and specific immunofluorescence labeling of samples prepared using a protocol similar that to prepare samples for analysis by electron microscopy (brain samples fixed in FA and glutaraldehyde and embedded in Lowicryl plastic). A total of 17 positives from the CBS evaluation of samples prepared in LR White plastic were tested on AT sections of mouse neocortex fixed in FA and glutaraldehyde and embedded in Lowicryl plastic; of these, 12 (71%) were positive (**Supplemental Table 1**).

Five of these mAbs was further evaluated at the EM level using immunogold labeling (**Figure 8**). The performance of the antibodies was considered to be good if they labeled the expected target with a high ratio of signal to noise. Both of the Homer mAbs identified using the AT CBS assay (L113/13 and L113/130, **Figures 6 and 7**) performed well when used for immunogold EM on Lowicryl HM20-embeded sections from mouse brain (**Figure 8A, B** and **Figures 8-1 and 8-2**). Three different mAbs against gephyrin (a postsynaptic protein at inhibitory synapses) that were similarly identified using the AT CBS assay (**Figure 8C-F**), were also confirmed as suitable for immunogold EM (**Figure 8G, H**). Thus, the AT CBS assay of LR White embedded samples also has a high predictive value for identifying Abs effective for postembedding immunogold electron EM on brain tissue embedded in Lowicryl HM20.

**Figure 8.**
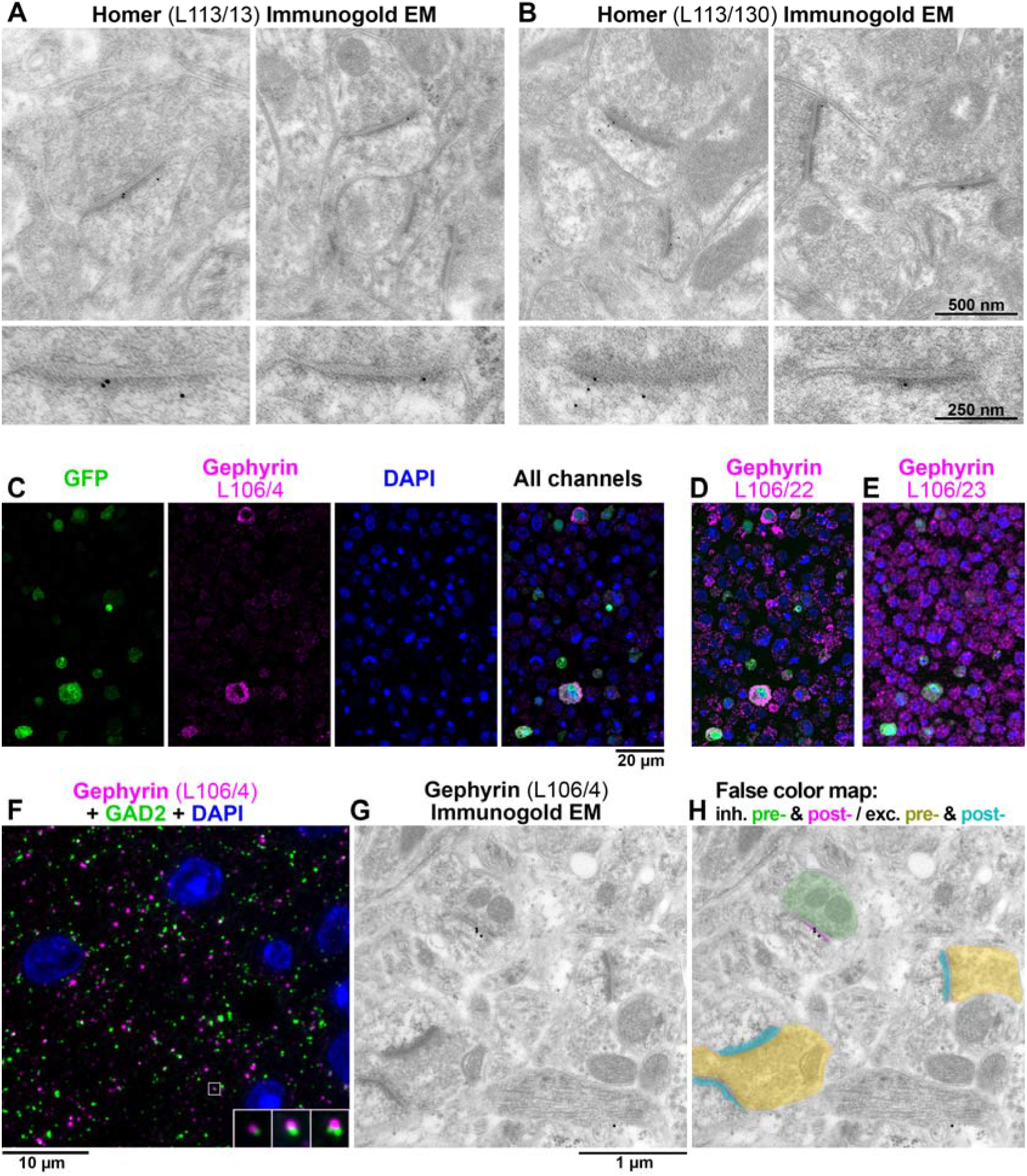
Ab screening for immunogold electron microscopy. **A, B.** Immunogold EM with L113/13 (A) and L113/30 (B) mAbs on Lowicryl HM20 embedded tissue from mouse cortex. The immunogold label localizes at asymmetric postsynaptic densities. The bottom panels show a magnified and rotated view of a postsynaptic density from the top panel. **C.** AT CBS assay for gephyrin mAb L106/4 immunolabeling of a 400 nm section from COS-1 cells co-transfected with separate plasmids encoding gephyrin and EGFP and embedded in LR White. The gephyrin immunolabeling colocalizes at the cellular level with EGFP marker expression. Because in this case the COS-1 cells were co-transfected with two separate plasmids, the overlap is not complete and some GFP-positive cells do not label with L106/4. **D.** An adjacent 400 nm section immunolabeled with mAb L106/22. While this mAb recognizes the transfected cells, there is also a high level of non-specific labeling and therefore it was rejected. **E.** mAb L106/23 does not label the transfected cells and was also rejected. **F.** AT immunofluorescence of an LR White-embedded 70 nm section from adult mouse cortex with L106/4 mAb against gephyrin (magenta), rabbit mAb GAD2 (Cell Signaling #5843, green) and DAPI (blue). The insert shows three consecutive sections through the synapse that is marked with a white box. **G**. Immunogold EM using the same L106/4 mAb on Lowicryl HM20 embedded tissue from mouse cortex. **H.** False color map of the section in G. The immunogold is associated with the postsynaptic side of the inhibitory synapse, but not excitatory synapses in the same field of view. See Supplementary Figures 1 and 2 for additional examples of Immunogold EM labeling.

### Application of the AT CBS assay to development of novel mAbs

We have completed 15 separate mAb projects, each targeting a distinct protein in which we used the CBS proxy assay to identify candidate mAbs that are effective for AT-based imaging. More than 1900 samples were screened with the CBS assay, and 259 CBS positive parents (CBS score ≥2) were identified. Out of the CBS positive parents, 207 were subsequently tested for AT on brain sections, and 124 out of these (60%) were also positive for AT on brain sections. Compared to the other assays that we performed, the CBS assay had a higher predictive value for identifying candidate mAbs positive for AT on brain sections (**Figure 9**). Thus, for the three projects (L113 Homer1, L109 Calbindin and L106 Gephyrin) where all top ELISA positive candidates (**Figure 3**) were selected for screening on every assay, the CBS screen had a higher positive predictive value for mAbs suitable for brain AT than any other assay, as well as a lower false omission rate. Positive predictive values were calculated as the number of candidates that were positive by both CBS assay and on brain AT (true positives), as a percent of all positive CBS candidates. False omission rate was calculated as the number of candidates negative on the CBS assay but positive for brain AT, as a percent of all the negative CBS candidates, and therefore reflects the likelihood of missing positive brain AT candidates (**Figure 9**).

**Figure 9.**
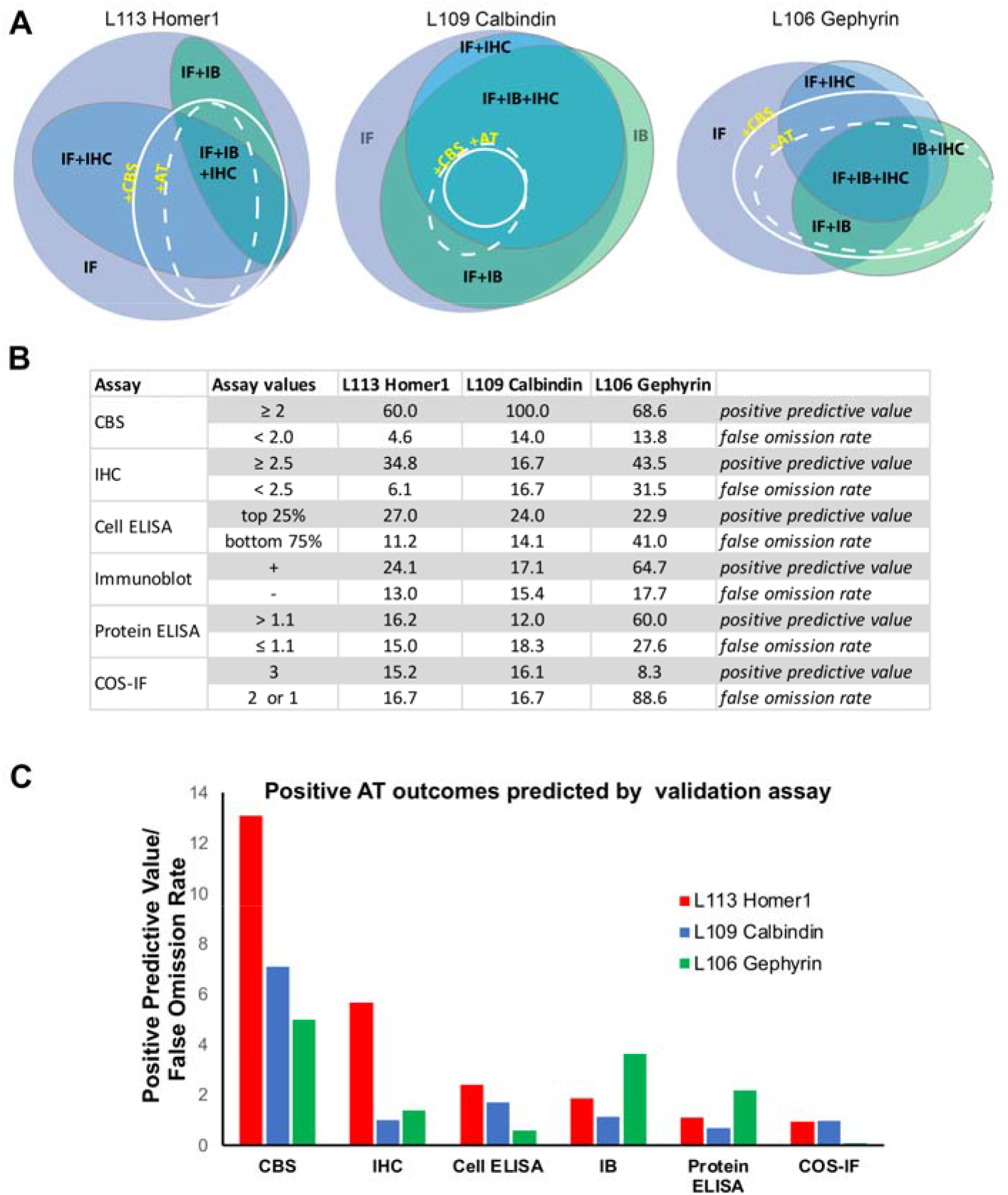
The CBS assay has high predictive value for Ab success in AT experiments. **A.** Euler diagrams for projects L113 (Homer1), L109 (Calbindin) and L106 (Gephyrin). **B** Table listing the percent of ELISA positive candidates giving rise to brain section AT positive mAbs (score ≥ 2.5) broken down by their performance on each validation assay. **C.** Ratio of the positive predictive value for AT suitable mAbs to the false omission rate shown for each validation assay. The CBS assay was most predictive for identifying brain AT positive mAbs.

Overall, from the 15 projects, 12 yielded AT-validated mAbs (**Supplemental Table 1**). Two projects did not yield any positive candidates on the AT CBS screen, although conventional IHC positives were obtained from both projects. Another project (L125 targeting synapsin-3) failed for reasons unrelated to AT, as all of the obtained candidate mAbs were found to exhibit cross-reactivity to synapsin-1. To verify that we were not inadvertently excluding candidate mAbs with potential AT utility by applying the AT CBS as a filter, for the two projects that did not yield any AT CBS positive candidates we also tested a set of candidate mAbs that had high scores from the IHC screen, but none of these yielded labeling in the AT brain assay.

For many projects the CBS assay enabled us to develop more than one AT-validated mAb against the same target protein, either yielding mAbs of different mouse IgG subclasses (L106, L109, L122), or different target isoform specificity (L113, L127). These results support the reliability of the AT CBS assay for identifying a subpopulation of Abs that have a high likelihood of exhibiting labeling of their target in AT brain sections. The overall list of Abs tested and results can be found in **Supplemental Table 1**.

## Discussion

The lack of highly validated Abs for research is a widely-recognized problem (Begley and Ellis, 2012; Baker, 2015a, b) that has forced laboratories to employ extensive in-house Ab testing prior to their use (Bordeaux et al., 2010; Micheva et al., 2010a). Here we describe a systematic, rapid, and effective approach to validate Abs for brain AT, leading to a robust set of mAbs available to the research community (**Supplemental Table 2**). We introduce a simple and low-cost proxy assay with a high predictive value for Abs effective and specific for immunolabeling of AT brain sections. Unlike direct screening on AT brain samples, the cell-based proxy screen does not use samples from animals, reducing animal use. The visual analysis of the CBS assay is also more straightforward than AT screening. It utilizes a heterologous expression system, allowing specific candidate Ab labeling of transfected cells to be easily distinguished from non-specific labeling of neighboring non-transfected cells. Visual comparison of transfected to non-transfected cells is much simpler than evaluation of AT brain samples, which must be performed at the level of individual synapses and may be confounded by synapse variability, low levels of expression or unknown distribution. Finally, because target proteins are overexpressed in transfected heterologous cells, the CBS assay is more sensitive at detecting Abs that may be of low concentration in hybridoma supernatants early in the development pipeline. This is particularly advantageous as the mAb candidate screening employs supernatants from hybridoma cultures, often with low levels of Ab, at a time in the development workflow when emphasis is on maintaining the health of the hybridoma cells prior to their cryopreservation, not on maximizing mAb accumulation in the medium. Moreover, at this stage hybridomas are not typically monoclonal; accordingly, these supernatants can contain multiple representations of target specific mAb at very different concentrations, which may be lower than typically present in final mAb preparations after subcloning to monoclonality and growing under culture conditions designed to yield maximal mAb accumulation in the medium. Therefore, an elevated level of target protein expression facilitates successful labeling during early stage screening.

In the experiments reported here, we included AT CBS screens in the workflow for 15 different projects, each aimed at developing mAbs against a distinct target. Each project employed screens aimed at developing mAbs for use in multiple downstream applications including transfected cell immunocytochemistry, brain immunoblots, immunohistochemistry on FA fixed conventional brain sections, and brain AT (**Figures 4 and 5**). Comparing the outcomes with Euler diagrams (Micallef and Rodgers, 2014) (https://www.eulerdiagrams.org/eulerAPE/), shows that performance in one of these applications does not predict success in other applications (**Figure 9**), highlighting the need for application-specific screening. This confirmed our experience with many commercially available Abs, which were validated in applications other than AT, and often failed when used for AT.

Our results suggest that the false omission rate of the AT CBS assay relative to brain AT is quite low (≈5-15%). However, the AT CBS assay does yield “false positives” (candidate mAbs that work in the AT CBS assay but not for brain AT), likely because the AT CBS assays involve overexpression of the target protein in the transfected cells. Moreover, the use of heterologous cells means that the target protein may not undergo the same posttranslational modifications as it does in neurons, and proteins that interact with the target protein in neurons may not be expressed; accordingly, epitopes targeting posttranslational modifications or protein-protein interactions may be accessible in heterologous cells but not in brain samples. This is especially problematic for synapse-associated proteins, which are extensively regulated by posttranslational modifications and participate in complex and densely packed networks of interacting proteins (Grant, 2006; Zhu et al., 2018; Grant, 2019; Helm et al., 2021). This dense protein network can also result in more general Ab access problems, since access of the Ab to the synaptic compartment may be limited. Ineffective immunolabeling of synaptic targets due to these considerations led to the development of numerous protocols for enhancing Ab access in conventionally prepared brain sections (Watanabe et al., 1998; Fukaya and Watanabe, 2000). While ultrathin samples such as those prepared for AT are expected to have fewer issues with macro level Ab access to the synaptic compartment, there may still remain access problems at the molecular level, and these samples would presumably retain fixative-stabilized protein-protein interactions not present in heterologous cells, resulting in ineffective labeling of occluded epitopes at synapses but not in heterologous cells. Despite the occasional false-positives, the proxy CBS assay was effective at filtering out the numerous candidates that score as negatives in both cell-and brain -based AT. In the exemplar Homer1 project there was a marked increase in the success rate in brain AT evaluation for those judged positive in the CBS assay (21/35 ≈ 60% of candidates with brain AT scores >2.5) compared to the success rate for the overall pool of 144 candidates (22/144 ≈ 15% of candidates with brain AT scores >2.5) that would have required evaluation had the CBS assay filter not been employed. Moreover, there was an extremely low false omission rate in the CBS assay (5 candidates with a brain AT score >2.5 out of 109 candidates with CBS score ≤1.9). In contrast, other standard assays (conventional immunocytochemistry against transfected heterologous cells, immunoblots, conventional immunohistochemistry against brain sections) lacked predictive value for AT-effective Abs (**Figure 9**).

Our results suggest that the same principles for Ab screening can be applied to other postembedding brain immunolabeling applications, including immunogold EM. In recent years, large-scale volume EM has provided important insights into the microscale organization of brain and principles of neuron connectivity (Zheng et al., 2018; Scheffer et al., 2020; Bae et al., 2021; Shapson-Coe et al., 2021). Complementing such expansive ultrastructural data with molecular information is rare (Anderson et al., 2011; Shahidi et al., 2015), due in large part to the lack of Abs suitable for postembedding immunoelectron microscopy. Our experiments suggest that the CBS proxy assay can also be used to identify Abs with high probability of success for immunogold EM, thus providing an efficient preliminary screen for suitable reagents for this highly demanding and resource-intensive application. Expanding the repertoire of synaptic Abs for electron microscopy applications will further increase the ability to collect correlated molecular and ultrastructural data in future connectomics studies. We suggest that this assay strategy could be employed whenever substantial collections of Abs against a given target need to be evaluated for AT or immunoelectron microscopy.

## Acknowledgements

We thank Camelia Dumitras, James Terry, Nafisa Ghori and Jenna Schardt for excellent technical assistance.

## Supplemental Legends

**Supplemental Table 1**. **Table of all antibodies tested for AT.** RRID, Research Resource Identifier. “Tested In” column shows the species where the antibody was tested. M, mouse, H, human, R, rat, Z, zebrafish. The results from the antibody testing indicated in the next column, Target Label, apply to all species where the antibody was tested, unless explicitly stated. Testing was performed on formaldehyde fixed tissue embedded in LR White, except for the antibodies indicated with * which require glutaraldehyde in the fixative. The last column, Performance on Lowicryl, indicates results of the testing on tissue fixed with a combination of formaldehyde and glutaraldehyde and embedded in Lowicryl HM20.

**Supplemental Table 2. Commonly used antibodies for AT, grouped by target.** These antibodies have been validated in both mouse and human biopsy brain tissue, except where indicated (**). The great majority of antibodies perform well with both formaldehyde fixation and with a combination of formaldehyde and glutaraldehyde, except several that require glutaraldehyde in the fixative (*). RRID, Research Resource Identifier.

**Supplemental Figure 1: L113/13 immunogold EM on mouse tissue.** Examples of L113/13 immunogold labeled synapses from mouse neocortex embedded in Lowicryl HM20.

**Supplemental Figure 2: L113/130 immunogold EM on mouse tissue.** Examples of L113/130 immunogold labeled synapses from mouse neocortex embedded in Lowicryl HM20.

## Bibliography

Anderson JR, Jones BW, Watt CB, Shaw MV, Yang JH, Demill D, Lauritzen JS, Lin Y, Rapp KD, Mastronarde D, Koshevoy P, Grimm B, Tasdizen T, Whitaker R, Marc RE (2011) Exploring the retinal connectome. Mol vis 17:355–379.

Bae JA et al. (2021) Functional connectomics spanning multiple areas of mouse visual cortex. bioRxiv: 2021.2007.2028.454025.

Baker M (2015a) Antibody anarchy: A call to order. Nature 527:545–551.

Baker M (2015b) Reproducibility crisis: Blame it on the antibodies. Nature 521:274–276.

Begley CG, Ellis LM (2012) Drug development: Raise standards for preclinical cancer research. Nature 483:531–533.

Bekele-Arcuri Z, Matos MF, Manganas L, Strassle BW, Monaghan MM, Rhodes KJ, Trimmer JS (1996) Generation and characterization of subtype-specific monoclonal antibodies to K+ channel alpha-and beta-subunit polypeptides. Neuropharmacology 35:851–865.

Boiko T, Rasband MN, Levinson SR, Caldwell JH, Mandel G, Trimmer JS, Matthews G (2001) Compact myelin dictates the differential targeting of two sodium channel isoforms in the same axon. Neuron 30:91–104.

Bordeaux J, Welsh A, Agarwal S, Killiam E, Baquero M, Hanna J, Anagnostou V, Rimm D (2010) Antibody validation. BioTechniques 48:197–209.

Brorson SH. (2001) Deplasticizing or etching of epoxy sections with different concentrations of sodium ethoxide to enhance the immunogold labeling. Micron. 32:101–105.

Brandstatter JH, Dick O, Boeckers TM (2004) The postsynaptic scaffold proteins ProSAP1/Shank2 and Homer1 are associated with glutamate receptor complexes at rat retinal synapses. J Comp Neurol 475:551–563.

Collman F, Buchanan J, Phend KD, Micheva KD, Weinberg RJ, Smith SJ (2015) Mapping synapses by conjugate light-electron array tomography. J Neurosci 35:5792–5807.

Fagni L, Worley PF, Ango F (2002) Homer as both a scaffold and transduction molecule. Sci STKE 2002:re8.

Fukaya M, Watanabe M (2000) Improved immunohistochemical detection of postsynaptically located PSD-95/SAP90 protein family by protease section pretreatment: a study in the adult mouse brain. J Comp Neurol 426:572–586.

Gong B, Murray KD, Trimmer JS (2016) Developing high-quality mouse monoclonal antibodies for neuroscience research - approaches, perspectives and opportunities. N Biotechnol 33:551–564.

Grant SG (2006) The synapse proteome and phosphoproteome: a new paradigm for synapse biology. Biochem Soc Trans 34:59–63.

Grant SGN (2019) Synapse diversity and synaptome architecture in human genetic disorders. Hum Mol Genet 28:R219–R225.

Helm MS, Dankovich TM, Mandad S, Rammner B, Jahne S, Salimi V, Koerbs C, Leibrandt R, Urlaub H, Schikorski T, Rizzoli SO (2021) A large-scale nanoscopy and biochemistry analysis of postsynaptic dendritic spines. Nat Neurosci 24:1151–1162.

Holderith N, Heredi J, Kis V, Nusser Z (2020) A High-Resolution Method for Quantitative Molecular Analysis of Functionally Characterized Individual Synapses. Cell Rep 32:107968.

Lorincz A, Nusser Z (2008) Specificity of immunoreactions: the importance of testing specificity in each method. J Neurosci 28:9083–9086.

Meyer MP, Trimmer JS, Gilthorpe JD, Smith SJ (2005) Characterization of zebrafish PSD-95 gene family members. J Neurobiol 63:91–105.

Micallef L, Rodgers P (2014) eulerAPE: drawing area-proportional 3-Venn diagrams using ellipses. PLoS One 9:e101717.

Micheva KD, Smith SJ (2007) Array tomography: a new tool for imaging the molecular architecture and ultrastructure of neural circuits. Neuron 55:25–36.

Micheva KD, Busse B, Weiler NC, O’Rourke N, Smith SJ (2010a) Single-synapse analysis of a diverse synapse population: proteomic imaging methods and markers. Neuron 68:639–653.

Micheva KD, O’Rourke N, Busse B, Smith SJ (2010b) Array tomography: high-resolution three-dimensional immunofluorescence. Cold Spring Harb Protoc 2010:pdb top89.

Micheva KD, Chang EF, Nana AL, Seeley WW, Ting JT, Cobbs C, Lein E, Smith SJ, Weinberg RJ, Madison DV (2018) Distinctive Structural and Molecular Features of Myelinated Inhibitory Axons in Human Neocortex. eNeuro 5.

O’Rourke NA, Weiler NC, Micheva KD, Smith SJ (2012) Deep molecular diversity of mammalian synapses: why it matters and how to measure it. Nat Rev 13:365–379.

Perez-Otano I, Schulteis CT, Contractor A, Lipton SA, Trimmer JS, Sucher NJ, Heinemann SF (2001) Assembly with the NR1 subunit is required for surface expression of NR3A-containing NMDA receptors. J Neurosci 21:1228–1237.

Rasband MN, Trimmer JS (2001) Developmental clustering of ion channels at and near the node of Ranvier. Dev Biol 236:5–16.

Rasband MN, Park EW, Vanderah TW, Lai J, Porreca F, Trimmer JS (2001) Distinct potassium channels on pain-sensing neurons. Proc Nat Acad Sci USA 98:13373–13378.

Rasband MN, Park EW, Zhen D, Arbuckle MI, Poliak S, Peles E, Grant SG, Trimmer JS (2002) Clustering of neuronal potassium channels is independent of their interaction with PSD-95. J Cell Biol 159:663–672.

Rhodes KJ, Trimmer JS (2006) Antibodies as valuable neuroscience research tools versus reagents of mass distraction. J Neurosci 26:8017–8020.

Rhodes KJ, Trimmer JS (2008) Antibody-based validation of CNS ion channel drug targets. J Gen Physiol 131:407–413.

Scheffer LK et al. (2020) A connectome and analysis of the adult Drosophila central brain. Elife 9.

Schindelin J, Arganda-Carreras I, Frise E, Kaynig V, Longair M, Pietzsch T, Preibisch S, Rueden C, Saalfeld S, Schmid B, Tinevez JY, White DJ, Hartenstein V, Eliceiri K, Tomancak P, Cardona A (2012) Fiji: an open-source platform for biological-image analysis. Nat Methods 9:676–682.

Schrand AM, Schlager JJ, Dai L, Hussain SM (2010) Preparation of cells for assessing ultrastructural localization of nanoparticles with transmission electron microscopy. Nat Protoc 5:744–757.

Shahidi R, Williams EA, Conzelmann M, Asadulina A, Veraszto C, Jasek S, Bezares-Calderon LA, Jekely G (2015) A serial multiplex immunogold labeling method for identifying peptidergic neurons in connectomes. Elife 4.

Shapson-Coe A et al. (2021) A connectomic study of a petascale fragment of human cerebral cortex. bioRxiv:2021.2005.2029.446289.

Shiraishi-Yamaguchi Y, Furuichi T (2007) The Homer family proteins. Genome Biol 8:206.

Simhal AK, Gong B, Trimmer JS, Weinberg RJ, Smith SJ, Sapiro G, Micheva KD (2018) A Computational Synaptic Antibody Characterization Tool for Array Tomography. Front Neuroanat 12:51.

Simhal AK, Aguerrebere C, Collman F, Vogelstein JT, Micheva KD, Weinberg RJ, Smith SJ, Sapiro G (2017) Probabilistic fluorescence-based synapse detection. PLoS Comput Biol 13:e1005493.

Thevenaz P, Ruttimann UE, Unser M (1998) A pyramid approach to subpixel registration based on intensity. IEEE Trans Image Process 7:27–41.

Tiffany AM, Manganas LN, Kim E, Hsueh YP, Sheng M, Trimmer JS (2000) PSD-95 and SAP97 exhibit distinct mechanisms for regulating K(+) channel surface expression and clustering. J Cell Biol 148:147–158.

Trimmer JS, Trowbridge IS, Vacquier VD (1985) Monoclonal antibody to a membrane glycoprotein inhibits the acrosome reaction and associated Ca2+ and H+ fluxes of sea urchin sperm. Cell 40:697–703.

Watanabe M, Fukaya M, Sakimura K, Manabe T, Mishina M, Inoue Y (1998) Selective scarcity of NMDA receptor channel subunits in the stratum lucidum (mossy fibre-recipient layer) of the mouse hippocampal CA3 subfield. Eur J Neurosci 10:478–487.

Xiao B, Tu JC, Worley PF (2000) Homer: a link between neural activity and glutamate receptor function. Curr Opin Neurobiol 10:370–374.

Zheng Z et al. (2018) A Complete Electron Microscopy Volume of the Brain of Adult Drosophila melanogaster. Cell 174:730–743 e722.

Zhu F, Cizeron M, Qiu Z, Benavides-Piccione R, Kopanitsa MV, Skene NG, Koniaris B, DeFelipe J, Fransen E, Komiyama NH, Grant SGN (2018) Architecture of the Mouse Brain Synaptome. Neuron 99:781–799 e710.

