## Supplemental Table 2 for "Developing a Toolbox of Antibodies Validated for Array Tomography-Based Imaging of Brain Synapses"

**Figure 1-2. Commonly used antibodies for array tomography.** These antibodies have been validated in both mouse and human biopsy brain tissue, except where indicated (**). The great majority of antibodies perform well with both formaldehyde fixation and with a combination of formaldehyde and glutaraldehyde, except several that require glutaraldehyde in the fixative (*). RRID, Research Resource Identifier.

| **Antigen** | **Host** | **Antibody Source** | **Dilution** | **RRID** |
| --- | --- | --- | --- | --- |
| ***Neurotransmitters*** | | | | |
| GABA* | Guinea pig | Millipore AB175 | 1:5000 | AB_91011 |
| GABA* | Rabbit | Millipore AB131 | 1:10000 | AB_2278931 |
| Glutamate* | Rabbit | Millipore AB5018 | 1:100 | AB_91640 |
| ***Synaptic*** | | | | |
| Synapsin 1 | Rabbit | Cell Signaling 5297 | 1:500 | AB_2616578 |
| Synapsin 1/2 | Chicken | Synaptic Systems 106 006 | 1:100 | AB_2622240 |
| Synaptophysin | Mouse | Abcam ab8049 | 1:50 | AB_2198854 |
| PSD95 | Rabbit | Cell Signaling 3450 | 1:200 | AB_2292883 |
| PSD95 | Mouse | NeuroMab 75-348, clone K28/74 | 1:100 | AB_2315909 |
| Homer1 | Mouse | NeuroMab 75-454, clone L113/130 | 1:50 | AB_2877198 |
| VGluT1 | Guinea pig | Millipore AB5905 | 1:5000 | AB_2301751 |
| VGluT2 | Rabbit | Synaptic systems 135 403 | 1:100 | AB_887883 |
| GAD2 | Rabbit | Cell Signaling 5843 | 1:200 | AB_10835855 |
| Gephyrin | Mouse | NeuroMab 75-465, clone L106/4 | 1:100 | AB_2716264 |
| Clathrin** | Rabbit | Cell Signaling 4796 | 1:100 | AB_10557412 |
| ***Cell type markers*** | | | | |
| Parvalbumin | Rabbit | SWANT PV28 | 1:300 | AB_10013386 |
| Calbindin** | Rabbit | Millipore AB1778 | 1:50 | AB_2068336 |
| Calbindin | Mouse | NeuroMab 75-448, clone L109/57 | 1:100 | AB_2686913 |
| NPY | Mouse | NeuroMab 75-456, clone L115/13 | 1:100 | AB_2877570 |
| nNOS | Mouse | NeuroMab 73-480, clone L121/25 | 1:2 | AB_2651159 |
| Tyrosine hydroxylase | Rabbit | Millipore AB152 | 1:200 | AB_390204 |
| ***Receptors*** | | | | |
| GluA1 | Rabbit | Millipore #AB1504 | 1:100 | AB_2113602 |
| GluA2 | Rabbit | Abcam ab206293 | 1:50 | AB_2800401 |
| GluA2/3 | Rabbit | Millipore AB1506 | 1:100 | AB_90710 |
| GluA3 | Mouse | Millipore MAB5416 | 1:1000 | AB_2113897 |
| GluA4 | Rabbit | Cell Signaling 8070 | 1:50 | AB_10829469 |
| GluN1 | Mouse | Synaptic systems 114 011 | 1:500 | AB_887750 |
| GluN2B | Mouse | NeuroMab 75-101, clone N59/36 | 1:500 | AB_2232584 |
| GABA A Receptor β2/3** | Mouse | Millipore MAB341 | 1:100 | AB_2109419 |
| ***Cytoskeletal*** | | | | |
| α Tubulin | Rabbit | Abcam ab18251 | 1:200 | AB_2210057 |
| Acetylated α tubulin | Mouse | Sigma T6793 | 1:100 | AB_477585 |
| βIII Tubulin** | Chicken | Abcam ab41489 | 1:100 | AB_727049 |
| Detyrosinated tubulin** | Rabbit | Millipore AB3201 | 1:100 | AB_177350 |
| δ2 Tubulin** | Rabbit | Millipore AB3203 | 1:100 | AB_177351 |
| Neurofilament 200 | Rabbit | Sigma N4142 | 1:100 | AB_477272 |
| Neurofilament, heavy chain | Chicken | AVES NF-H | 1:100 | AB_2313552 |
| Neurofilament, light chain** | Chicken | AVES NF-L | 1:100 | AB_2313553 |
| γ actin | Mouse | Sigma A8481 | 1:100 | AB_2289264 |
| Synaptopodin** | Rabbit | Synaptic Systems 163 002 | 1:500 | AB_887825 |
| ***Glial*** | | | | |
| Glutamine Synthetase | Mouse | BD Biosciences 610517 | 1:25 | AB_397879 |
| GFAP | Chicken | AVES GFAP | 1:100 | AB_2313547 |
| ***Myelin*** | | | | |
| MBP | Chicken | AVES MBP | 1:100 | AB_2313550 |
| PLP | Chicken | AVES PLP | 1:100 | AB_2313560 |
| CNPase | Chicken | AVES CNP | 1:100 | AB_2313538 |
| ***Ion channels*** | | | | |
| Kv1.1** | Mouse | NeuroMab 75-105, clone K36/15 | 1:100 | AB_2128566 |
| Kv1.2** | Mouse | NeuroMab 75-008, clone K14/16 | 1:100 | AB_2296313 |
| Kv2.1** | Mouse | NeuroMab 75-014, clone K89/34 | 1:100 | AB_10673392 |
| ***Mitochondria*** | | | | |
| MDH2 | Rabbit | Origene TA308153 | 1:200 | AB_2722674 |
| TOMM20 | Rabbit | Abcam ab78547 | 1:100 | AB_2043078 |
| VDAC | Rabbit | ProteinTech 10866-1-AP | 1:50 | AB_2257153 |
| ***Other*** | | | | |
| Laminin** | Rabbit | Sigma L9393 | 1:25 | AB_477163 |
| Caspr** | Mouse | NeuroMab 75-001, clone K65/35 | 1:100 | AB_2083496 |
| GFP | Chicken | GeneTex GTX13970 | 1:100 | AB_371416 |
| GFP | Mouse | NeuroMab 75-131, clone N86/8 | 1:100 | AB_10671445 |

*Require glutaraldehyde in the fixative, **Not tested in human
