## Supplementary figures and images for "Developing a Toolbox of Antibodies Validated for Array Tomography-Based Imaging of Brain Synapses"

### Supplemental Figure 1

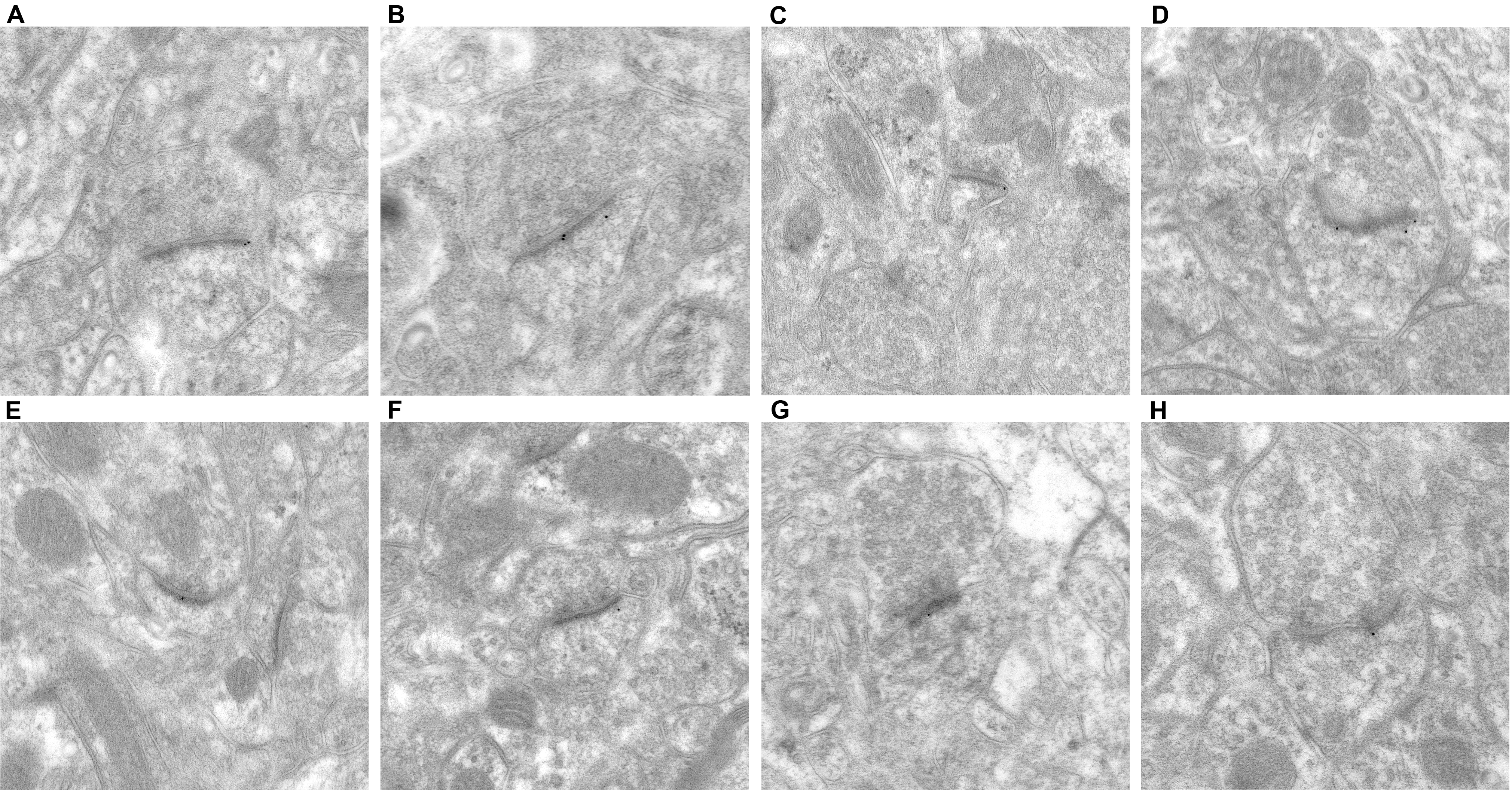

### Supplemental Figure 2

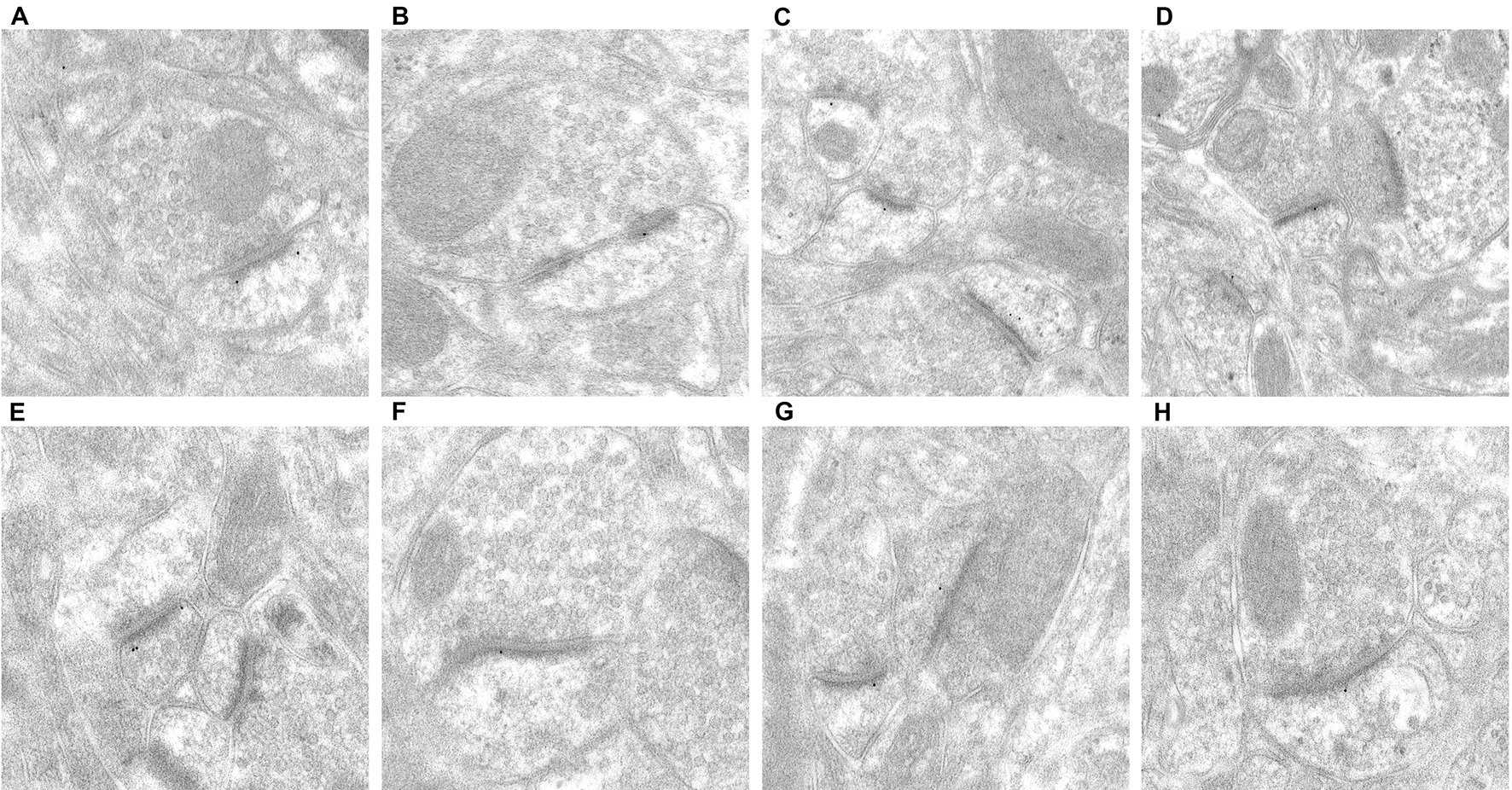
